# mTORC1 supports progression toward activation competence in quiescent adult neural stem cells

**DOI:** 10.64898/2026.05.04.722648

**Authors:** M. Thetiot, L. Taing, D. Morizet, G. Letort, L. Bally-Cuif

## Abstract

Neural stem cells (NSCs) sustain lifelong neurogenesis through the tight regulation of quiescence, self-renewal and differentiation. Quiescent NSCs (qNSCs) exist in distinct substates, ranging from deep to shallow quiescence, yet the mechanisms governing these transitions remain unclear. In long-term self-renewing NSCs of the adult zebrafish pallium, we show that mTORC1 activity is specifically enriched during a prolonged quiescence phase in which NSCs acquire activation competence. Functional perturbations, analyzed in situ and using single-cell RNA sequencing, reveal that mTORC1 regulates cell progression during this phase, concomitantly ensuring the correct tempo for NSC transition towards activation and the preservation of stemness. These findings challenge the classical view of mTORC1 as a simple regulator of proliferation and identify it as a key regulator of NSC quiescence heterogeneity and dynamics under physiological conditions. By coordinating stemness maintenance with activation competence, mTORC1 emerges as a central player balancing long-term NSC preservation with neurogenic output in the adult brain.

## INTRODUCTION

Adult stem cells (SCs), play a critical role in tissue maintenance and regeneration by balancing long-term self-renewal with the production of differentiated progeny. To preserve this potential over time, many SCs enter a reversible state of cell cycle arrest known as quiescence (usually G0). Quiescent SCs are not homogeneous: single-cell RNA sequencing (scRNA-seq), clonal lineage tracing and functional assays have revealed that SCs can occupy several molecular and functional substates ranging from “deep” to “shallow” quiescence. These substates differ in metabolic activity, activation competence - defined as the cells’ increased likelihood and responsiveness to enter activation upon stimulation- and lineage output (Belenguer et al. 2021; Mancini et al. 2023; Morizet et al. 2024; Dulken et al. 2017; Llorens-Bobadilla et al. 2015; Rigo et al. 2025; Rodgers et al. 2014; Laurenti et al. 2015; Rodgers et al. 2017), yet the intrinsic mechanisms that position SCs within these substates remain poorly understood. A central unresolved question is how SCs integrate internal clocks, including growth programs, with extrinsic cues to regulate their readiness to activate without compromising long-term maintenance.

The adult zebrafish pallium hosts a lifelong neurogenic niche, made of radial glial NSCs expressing markers such as glial fibrillary acidic protein (Gfap), glutamine synthetase (Gs) or the transcription factor Sox2 (Ganz et al. 2015; März et al. 2010; Suh et al. 2007). These NSCs can remain in quiescence for extended periods (weeks to months), a state promoted by Notch3 signaling (Alunni et al. 2013; Morizet et al. 2025). Upon activation, NSCs enter the cell cycle and can give rise to further committed neural progenitors (NPs) and ultimately neurons (Foley et al. 2024). NSCs and NPs (collectively termed NSPCs) form an epithelial monolayer whose apical surface lines the everted pallial ventricle. This organization enables long-term intravital imaging and the combined analysis of NSC states, fate decisions, and apical cellular features during neurogenesis (Mancini et al. 2023; Than-Trong et al. 2020). In transgenic Tg(*deltaA*:*Gfp*) animals reporting expression of the Notch ligand *deltaA*, we previously showed that the *deltaA*:Gfp^neg^ subset of pallial NSCs constitutes a self-renewing pool that undergoes asymmetric divisions to produce one *deltaA*:Gfp^neg^ NSC and one *deltaA*:Gfp^pos^ NSC biased toward neurogenesis (Mancini et al. 2023). We further showed that, post-division, quiescent self-renewing NSCs gradually enlarge their apical area (AA) over several months and that AA size positively correlates with their propensity to divide. However, self-renewing NSCs display marked heterogeneity in AA size and activation latency, and our results do not support the existence of defined size or temporal thresholds that would dictate activation and division. Rather, a slow cell-intrinsic process, possibly including metabolic or growth-related cues, might contribute to progressively shape NSC activation competence. The regulators of this intrinsic program remain unknown.

A mechanistic link between metabolic growth, quiescence depth, and activation readiness has been established in other stem cell systems. In muscle stem cells (MuSCs), the conserved serine-threonine kinase mTORC1, which integrates inputs from growth factors and nutrients to regulate cell growth and metabolism (Battaglioni et al. 2022; Shimobayashi et Hall 2014), drives the transition from deep quiescence to a metabolically primed “G_alert_” state upon injury, increasing cell size and responsiveness to activating cues without triggering immediate proliferation (Rodgers et al. 2014, 2017). Whether similar principles operate in adult NSCs and under physiological conditions remains unclear, in part because, in the brain, mTORC1 is typically associated with proliferating progenitors rather than quiescent stem cells (Hartman et al. 2013; Zhou et al. 2018; Baser et al. 2019). Hyperactivation of mTORC1 in neonatal NSCs from the mouse subventricular zone (SVZ) promotes proliferation and symmetric neurogenic divisions (Hartman et al. 2013). mTORC1 in adult NSCs is also necessary for division entry when quiescence signals are abrogated (Zhou et al. 2018). Finally, reduced mTORC1 activity during division biases adult SVZ progenitors toward neuronal differentiation (Baser et al. 2019; Carvajal Ibañez et al. 2023). These findings indicate that mTORC1 influences NSC activation and fate decisions at division. However, whether mTORC1 also operates during the quiescence phase has not been described.

While investigating quiescence heterogeneity, we unexpectedly detected mTORC1 activity in a measurable proportion of quiescent NSCs under physiological conditions in the adult zebrafish pallium. These findings prompted us to explore a potential new role for mTORC1 in NSC quiescence state dynamics. Combining pharmacological manipulations in vivo, 3D NSC spheroids, and single-cell transcriptomics upon mTORC1 inhibition, we show that mTORC1 engages sequentially along the NSC quiescence-to-activation trajectory: first marking a long phase of activation competence acquisition during quiescence, then rising during early activation prior to cell-cycle entry and finally becoming essential for successful cell-cycle progression. Importantly, mTORC1 inhibition in self-renewing NSCs not only arrests proliferation, as previously shown, but also delays and biases their progression along this continuum toward activation. Together, these findings identify mTORC1 as a key regulator of quiescence dynamics that drives the acquisition of activation competence and thereby balances NSC maintenance with neurogenic output in the adult brain.

## RESULTS

### mTORC1 activity reveals functional heterogeneity among adult pallial NSCs

To assess endogenous mTORC1 activity across adult pallial neural stem and progenitor cells (NSPCs) of the zebrafish brain, we monitored phosphorylation of the ribosomal protein S6 (pS6), a downstream effector of the mTORC1-S6K pathway that links nutrient-sensing signals to cell growth and proliferation (Shimobayashi et Hall 2014). Using whole-mount immunohistochemistry (IHC) on dorsal pallia from 3-month-old Tg(*gfap:Gfp)* (Bernardos et Raymond 2006) zebrafish, we labeled radial glial NSCs using Gfp (cells positive for Gfp will be referred to *gfap*^pos^), outlined apical NSPC contours with the tight junction protein Zona occludens 1 (Zo1), and detected mTORC1 activity with pS6. Analyses were restricted to the dorsal medial (Dm) pallial domain, a region of sustained neurogenesis (Dray et al. 2021), thereby ensuring spatial consistency and controlling for known regional heterogeneity across pallial subdivisions (Ganz et al. 2015; Pandey et al. 2023).

We observed cytoplasmic pS6 signal in a subset of NSPCs, with pS6 predominantly enriched in *gfap*^pos^ neural stem cells (NSCs), but rarely detected in *gfap*^neg^ neural progenitors (NPs) **(Figure 1a)**. A treatment with the mTORC1 inhibitor Rapamycin, applied in vivo for 24h on adult fish, strongly diminished pS6 signal **(Figure S1a)**, confirming that it results from mTORC1 activity. Using a custom image analysis pipeline combining progenitor segmentation with manual assignment (Letort et al. 2026), we quantified pS6 and *gfap*:Gfp expression. Approximately half of NSPCs (48.2 ± 2.8%) displayed detectable pS6 signal **(Figure S1b)**, and most of these pS6^pos^ cells corresponded to *gfap*^pos^ NSCs **(Figure 1b)**, indicating that mTORC1 activity is largely confined to the NSC population, where it is present in more than half of the cells at any time (57.1 ± 3.4%) **(Figure 1c)**. To determine whether pS6 expression reflects selective phosphorylation or differences in protein abundance, we examined total S6 levels. IHC for S6 revealed broad and uniform expression across NSPCs **(Figure S1c)**, indicating that differential pS6 labeling arises from post-translational regulation rather than differences in S6 availability. To corroborate these observations, we examined phosphorylation of 4EBP1 **(Figure S1d)**, another downstream target of mTORC1, and found a similar pattern: p4EBP1 was expressed in half of NSPCs **(Figure S1e)** and present in 49.3% of *gfap*^pos^ NSCs **(Figure S1f-g)**. Due to consistent overlap in expression patterns among NSCs, pS6 was used as the primary marker of mTORC1 activity throughout the study.

**Figure 1.**
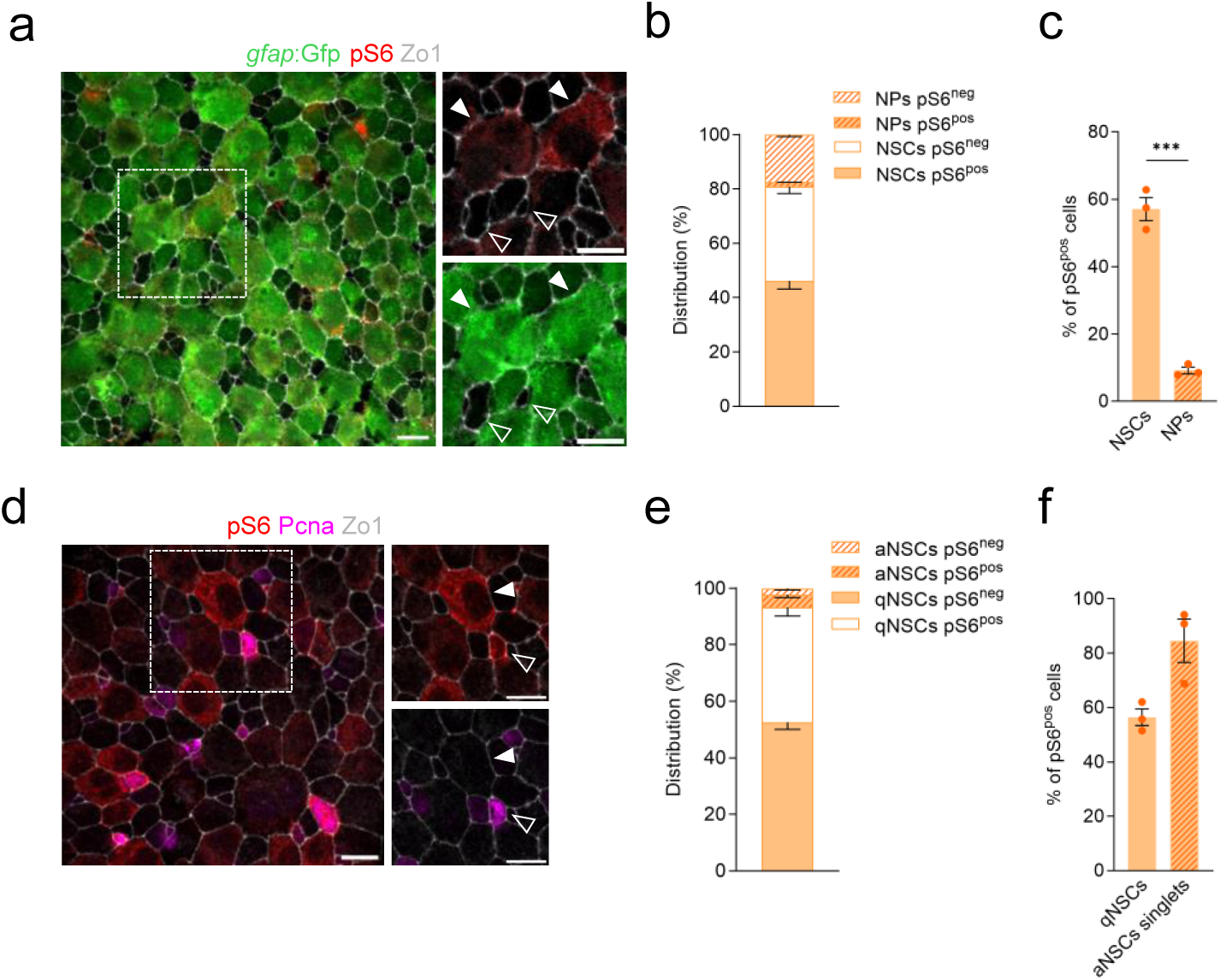
mTORC1 activity is enriched in a subset of adult zebrafish pallial NSCs. **(a)** Representative whole-mount apical view of the dorsal pallium (Dm) from 3-month-old *Tg(gfap:*Gfp) zebrafish processed in immunohistochemistry (IHC) for Gfp (green), pS6 (red), and Zo1 (white). Close-up views highlight examples of pS6^pos^/*gfap*^pos^ NSCs (white arrowheads) and pS6^neg^/gfap^neg^ NPs (empty arrowheads). **(b)** Distribution of pS6 expression among *gfap*^pos^ NSCs and *gfap*^neg^ NPs (full and stripped orange, respectively). **(c)** Percentage of pS6^pos^ cells among *gfap*^pos^ NSCs and *gfap*^neg^ NPs. **(d)** IHC on 3-month-old Tg(*gfap*:Gfp) adult pallia showing pS6 (red) expression in quiescent (Pcna^neg^, white arrowheads) and proliferating (Pcna^pos^, purple, empty arrowheads) NSPCs. **(e)** pS6 expression among quiescent (qNSCs) and activated (aNSCs) NSCs (full and stripped orange, respectively). **(f)** Percentage of pS6^pos^ qNSCs (left) and aNSCs pre-division (Pcna^pos^ singlets, right). The dashed white boxes localize high-magnification inserts. NSC: neural stem cell, NP: neural progenitor. NSPCs: neural stem and progenitor cells. Data shown as mean ± SEM from n=3 independent experiments. Two-tailed Mann–Whitney test. p-values : ****<0.0001; ***<0.001; **<0.01; *<0.05. Scale bars : 10 µm (a, d); inserts : 10 µm (a, d).

Activated NSCs have been previously defined by the expression of proliferation markers such as Ki67, Mcm2 or Pcna (Llorens-Bobadilla et al. 2015; Dulken et al. 2017; Morizet et al. 2025; Labusch et al. 2024). To interpret pS6 expression across NSC states, we used Pcna staining to separate quiescent NSCs (qNSCs, *gfap*^pos^/Pcna^neg^) and activated NSCs (aNSCs, *gfap*^pos^/Pcna^pos^) **(Figure 1d)**. Because Pcna marks the late G1/S/G2/M phases of the cell cycle, Pcna^pos^ cells were further classified into singlets (pre-division) and doublets (post-division), to resolve entry into proliferation. Most aNSCs displayed pS6 expression **(Figure 1e),** particularly Pcna^pos^ singlets (84.6 ± 8.0%) **(Figure 1f)**, consistent with the established role of mTORC1 in supporting proliferation. Surprisingly, however, more than half of qNSCs (56.4 ± 3.1%) also exhibited pS6 signal **(Figure 1e, 1f)**, indicating that mTORC1 activity is not restricted to proliferating progenitors but is present in a substantial fraction of qNSCs. These results suggest a previously undescribed role of mTORC1 during quiescence, in a cell substate- or subtype-specific manner.

### mTORC1 is required for NSC cell cycle entry in vitro

This unexpected observation prompted us to separately address mTORC1 functional requirements during NSC entry into proliferation and during the quiescent state. We first focused on the transition to activation **(Figure 2a)**. For this, we turned to a 3D in vitro culture system that we recently developed, which recapitulates key features of adult zebrafish pallial NSCs, including their ability to remain quiescent, proliferate and self-renew (Thetiot et al. 2025). NSPCs isolated from adult pallia were expanded in 2D-cultures to enrich for NSCs, then aggregated in 96-well plates where they self-organized into spheroids over several days. In the presence of mitogens, NSCs within these spheroids maintained expression of the stemness marker Sox2, and co-expressed *gfap*. Cultures could be maintained for up to a week, during which NSCs were found in both quiescent (qNSCs, *gfap*^pos^/Pcna^neg^) and activated (aNSCs, *gfap*^pos^/Pcna^pos^) states. By day 3, spheroids reached a stable equilibrium in which less than 10% of NSCs were proliferating under basal conditions. Thus, they provide a reproducible platform for controlled pharmacological manipulation, with a moderate enrichment for activation events compared to what is observed in vivo (**Figure 1e**).

**Figure 2.**
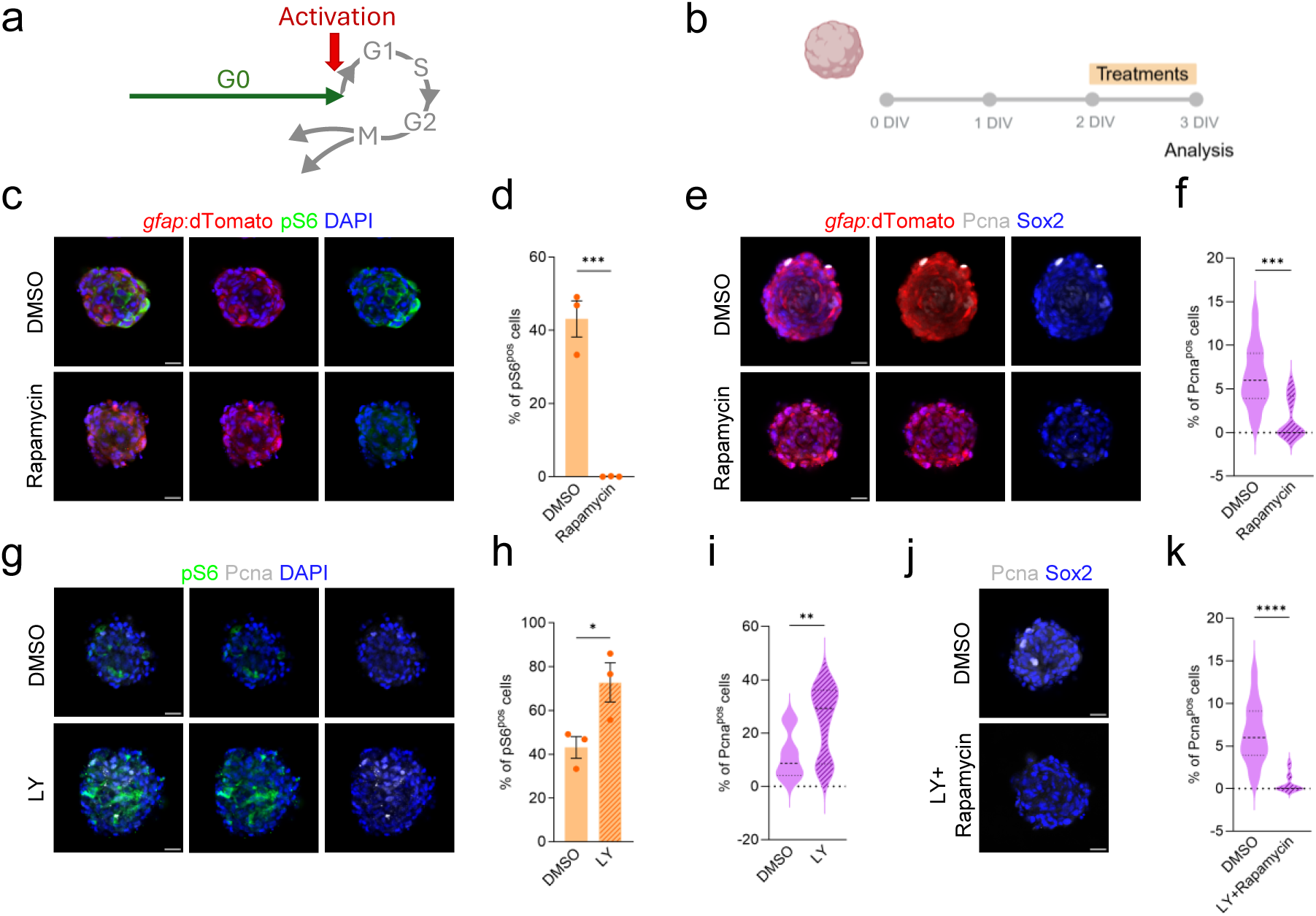
mTORC1 is required for NSC quiescence exit. **(a)** Positioning of the activation step as used in this paper. **(b)** Schematic representation of the treatment procedure. **(c)** Representative confocal images of 3D spheroids derived from Tg(*gfap*:dTomato) adult zebrafish NSCs treated for 24h with DMSO or Rapamycin, stained for pS6 (green), dTomato (red), and DAPI (blue). **(d)** Quantification of the percentage of pS6^pos^ NSCs (*gfap*^pos^) in DMSO and Rapamycin-treated spheroids after 24h. **(e)** Representative confocal images of 3D spheroids derived from Tg(*gfap*:dTomato) adult zebrafish NSCs treated for 24 h with DMSO or Rapamycin, stained for Pcna (green), dTomato (red), and Sox2 (blue). **(f)** Percentage of proliferating NSCs (*gfap*^pos^/Pcna^pos^) in control (DMSO) or Rapamycin-treated 3D spheroids after 24 h. **(g)** 3D spheroids treated for 24 h with DMSO or LY411575 (LY), stained for pS6 (green) and DAPI (blue). **(h)** Quantification of the percentage of pS6^pos^ cells in DMSO and LY-treated conditions (full and stripped orange, respectively). **(i)** Percentage of proliferating NSCs (*gfap*^pos^/Pcna^pos^) in control (DMSO) or LY-treated 3D spheroids after 24 h. **(j)** 3D spheres stained for Pcna (white) and Sox2 (blue) after 24h DMSO or LY+Rapamycin treatments. **(k)** Percentage of proliferating NSCs (*gfap*^pos^/Pcna^pos^) after 24h DMSO or LY+Rapamycin. Data shown as mean ± SEM from n=3 (d, f, h, i, k) independent experiments. Two-tailed non-parametric Mann–Whitney test (d, h) or unpaired t-test Welch correction in (f, i, k). p-values : ****<0.0001; ***<0.001; **<0.01; *<0.05. Scale bars : 20 µm (c, e, g, j).

To examine the functional requirement of mTORC1 on activation, spheroids generated from Tg(*gfap:*dTomato) NSCs were treated for 24h with DMSO (control) or the mTORC1 inhibitor Rapamycin **(Figure 2b)**. Immunostaining revealed that mTORC1 activity was robustly maintained under control conditions **(Figure 2c)** and, as observed in vivo, was present in both activated and quiescent NSCs, including 41.5 ± 6.3% of qNSCs **(Figure S2a)**. Total S6 protein levels were uniform across cells **(Figure S2b)**, confirming that the heterogeneity arises from phosphorylation and not protein abundance. Rapamycin efficiently abolished pS6, confirming pathway inhibition **(Figure 2c, 2d)** and significantly reduced the proportion of aNSCs (1.6 ± 0.6 vs 6.4 ± 0.9% in DMSO) **(Figure 2e, 2f)**, without affecting overall cell number (73.9 ± 13.9% (DMSO) vs 70.2 ± 12.5% (Rapamycin)) **(Figure S2c)**. This shows that mTORC1 activity contributes to NSC recruitment into the cell cycle.

To confirm this conclusion, we examined mTORC1 signaling during forced activation. In both zebrafish (Alunni et al. 2013; Morizet et al. 2025) and mouse (Zhang et al. 2019), Notch inhibition rapidly induces NSC re-entry into the cell cycle. 24h Notch inhibition with the γ-secretase inhibitor LY411575 (LY), applied to spheroids between days 2 and 3, increased the percentage of pS6-expressing NSCs (72.8 ± 9.0% vs 43.1 ± 4.9% in controls) **(Figure 2g, 2h)**, especially in qNSCs (64.6 ± 10.8% in LY-treated spheres vs 41.5 ± 6.2% in controls) **(Figure S2d)**, as well as the proportion of aNSCs (24.6 ± 3.3% vs 11.6 ± 2.1% in controls **(Figure 2i)**, indicating that mTORC1 upregulation is rapidly induced in activation. However, when LY and Rapamycin were applied simultaneous for 24h, proliferation remained strongly reduced (0.6 ± 0.3% vs 6.4 ± 0.9 in controls) **(Figure 2j, 2k)**, to levels indistinguishable from Rapamycin alone, despite the activation stimulus. Thus, preventing mTORC1 activity blocks the transition from quiescence to proliferation even under strong pro-activation conditions.

Together, these in vitro findings demonstrate that mTORC1 activity acts upstream of cell-cycle entry, enabling qNSCs to initiate the activation program and subsequent division. These findings are in line with previous work in adult mouse MuSCs and NSCs (Rodgers et al. 2014, 2017; Zhou et al. 2018; Hartman et al. 2013).

### mTORC1 is required for efficient NSC cell cycle progression under activation conditions in vivo

To assess mTORC1 function on activation and cell cycle progression in vivo, we treated adult fish with either DMSO or Rapamycin for 24h, 48h and 72h and analyzed the proliferation of pallial progenitors **(Figure S3a)**. Whole-mount IHC for Pcna and Zo1 revealed that the overall proportion of Pcna^pos^ cells upon Rapamycin treatment remained largely unchanged compared to controls (24h: 13.8 ± 2.1% vs 13.9 ± 1.4%; 48h: 10.3 ± 0.9 vs 9.1 ± 0.05%; 72h: 10.1 ± 2.6% vs 8.9 ± 4.6% in DMSO vs Rapamycin, respectively) **(Figure S3b, S3c)**. To refine this analysis, because self-renewing NSCs and neurogenic NSPCs display very different activation rates (average quiescence duration of ≈124 days for self-renewing NSCs compare to ≈28 days for neurogenic progenitors) (Mancini et al. 2023), we distinguished between these subpopulations using the Tg(*deltaA*:Gfp) background. As previously resolved by long-term live imaging (Mancini et al. 2023), *deltaA*:Gfp^neg^ NSCs constitute the long-term self-renewing pool, hierarchically upstream of *deltaA*:Gfp^pos^ cells, which include further committed NSPCs and are biased towards neurogenesis. Sensitive whole-mount IHC in adult Tg(*deltaA*:Gfp) pallia also revealed a smaller population of cells with weak *deltaA*:Gfp signal, undetectable in live observations but which share with *deltaA*:Gfp^neg^ NSCs their high expression of Gfap, size range and fate; in the following, we therefore refer to *delta*:Gfp^neg/weak^ as the self-renewing NSC population **(Figure 3a, 3b)**. Combining IHC for Gfp, Pcna and pS6, we confirmed that mTORC1 activity was prominent in most activated cells (self-renewing NSCs : 97.3 ± 3.7%, neurogenic NSPCs : 65.9 ± 11.7%) **(Figure 3c, 3d)**. Rapamycin treatment however remained without effect on the proportion of Pcna^pos^ cells even when NSC subtypes were resolved **(Figure S3d)** (self-renewing pool at 24h: 1.7 ± 0.7% vs 0.9 ± 0.4%; 48h: 0.6 ± 0.1 vs 0.8 ± 0.1%; 72h: 1.2 ± 0.9% vs 1.3 ± 1.1% in DMSO vs Rapamycin, respectively. Neurogenic pool at 24h: 27.0 ± 4.4% vs 27.6 ± 6.9%; 48h: 22.8 ± 1.0 vs 20.2 ± 1.0%; 72h: 24.7 ± 11.4% vs 17.2 ± 8.3% in DMSO vs Rapamycin, respectively).

**Figure 3.**
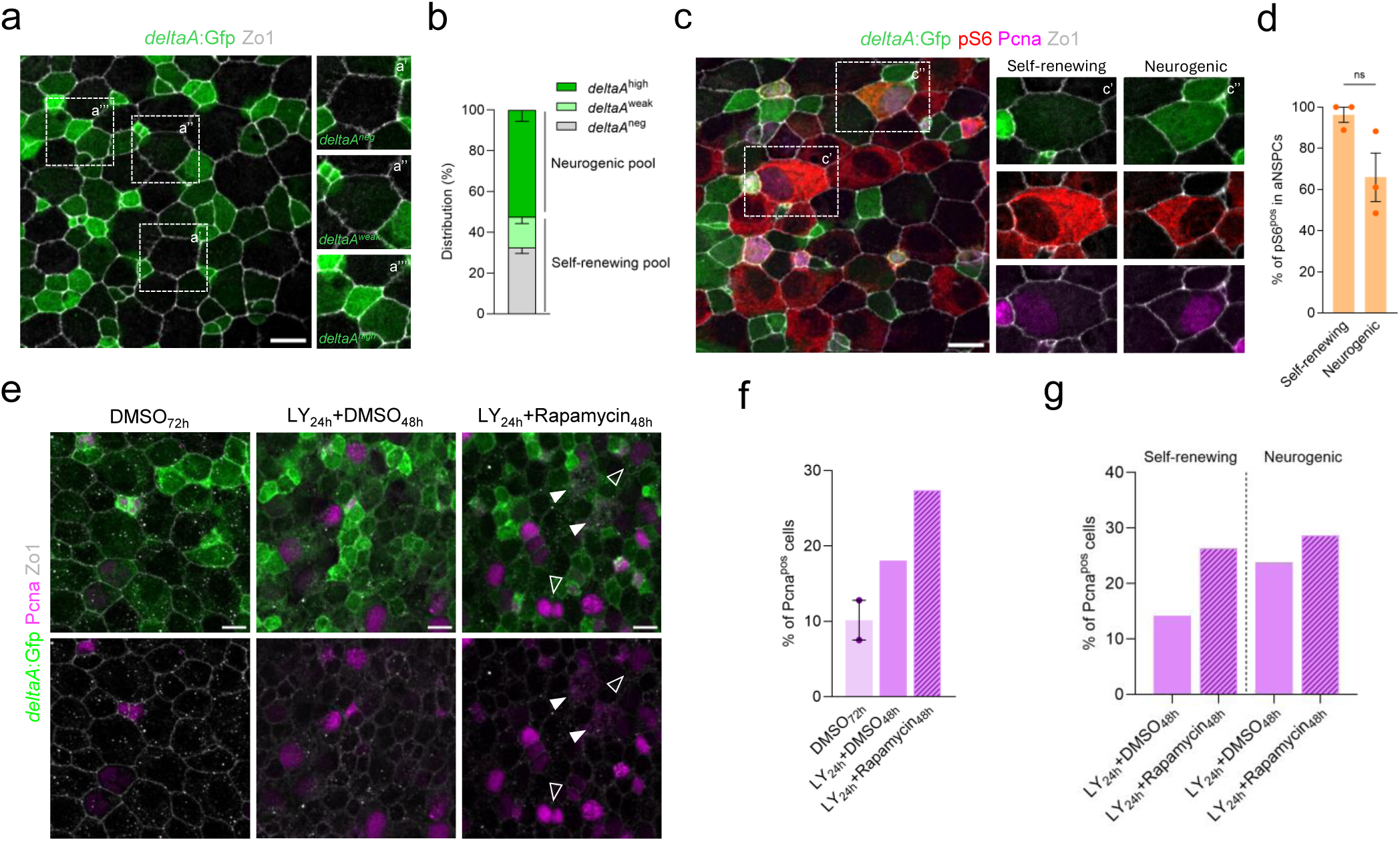
mTORC1 inhibition impairs NSC cell cycle completion following activation. **(a)** Whole-mount apical view of the pallium (Dm territory) in an adult Tg(*deltaA*:Gfp) fish stained for Gfp (green), and Zo1 (white), showing three categories of NSPCs based on *deltaA* expression levels : *deltaA*^neg^ (no Gfp) (a’), *deltaA*^weak^ (low Gfp) (a”) and *deltaA*^high^ (strong Gfp) (a”’). **(b)** Distribution of *deltaA*^neg/weak^ self-renewing and *deltaA*^high^ neurogenic NSPCs. Cells were manually classified based on Gfp intensity. **(c)** Whole-mount Dm apical view in an adult Tg(*deltaA*:Gfp) fish stained for pS6 (red), Pcna (magenta), Gfp (green), and Zo1 (white), highlighting self-renewing and neurogenic NSPC populations. **(d)** Percentage of pS6^pos^ among self-renewing and neurogenic aNSPCs (Pcna^pos^ singlets). **(e)** Representative images of whole-mount IHC from Tg(*deltaA*:Gfp) pallia showing Gfp (green), Pcna (magenta), and Zo1 (white) staining after the following treatments: DMSO (72h), LY (24h) followed by DMSO (48h), or LY (24h) followed by Rapamycin (48h). Nuclear abnormalities (e.g., enlarged nuclei and multinucleated (empty arrowheads), irregular chromatin (white arrowheads)) are present in pallial progenitors after LY_24h_+Rapamycin_48h_ treatment. **(f)** Percentage of Pcna^pos^ NSPCs in the whole population and **(g)** across *deltaA* subtypes (self-renewing *deltaA*^neg/weak^ and neurogenic *deltaA*^high^) after LY followed by DMSO or Rapamycin. The dashed white boxes localize high-magnification inserts. NSPCs: neural stem and progenitor cells. Data shown as mean ± SEM from n=3 (b, d) and n=1-2 (f, g) independent experiments. Two-tailed Mann–Whitney test. p-values : ****<0.0001; ***<0.001; **<0.01; *<0.05. Scale bars : 10 µm (a, c, e).

Given the intrinsically slow proliferation dynamics of adult NSCs in vivo, many of which remain quiescent for weeks (Mancini et al. 2023; Dray et al. 2021; Than-Trong et al. 2020), detecting significant changes in proliferation over short treatment windows was expectedly challenging. To overcome this limitation, and based on our in vitro results, we used a two-step activation paradigm that transiently pushes NSC towards activation before assessing the requirement of mTORC1 during cell cycle progression. We therefore treated Tg(*deltaA*:Gfp) adult fish with LY for 24h, followed by 48h treatment with either DMSO or Rapamycin. Treated fish were analyzed by IHC for Gfp, Pcna and Zo1 **(Figure 3e)**. As expected, LY_24h_+DMSO_48h_ led to a robust NSPC recruitment into proliferation among all NSPCs compared to DMSO_72h_ alone. LY_24h_+Rapamycin_48h_ resulted in a further increase in the overall proportion of Pcna^pos^ cells relative to controls **(Figure 3f)**. Notably, while *deltaA*^high^ had similar proportion of Pcna^pos^ between conditions, *deltaA*^neg/weak^ self-renewing NSCs displayed a significant accumulation of Pcna^pos^ cells compared to controls **(Figure 3g)**. In addition, self-renewing NSCs exhibited morphological signs of cell cycle disruption **(Figure 3e)**, representing about 13.2% of Pcna^pos^ NSCs from that pool. Thus, in vivo under our experimental conditions, Rapamycin does not prevent activation but disrupts orderly cell-cycle progression, leading to an accumulation of partially activated, cell cycle-stalled NSCs, a particularly prominent feature in self-renewing NSCs.

### mTORC1 activity during NSC quiescence is prominent in the self-renewing NSC pool

We next focused on mTORC1 activity during quiescence. To determine whether specific NSC subpopulations preferentially engage mTORC1 during quiescence, we examined the distribution of mTORC1 activity along lineage hierarchy using *deltaA*:Gfp expression, focusing on Pcna^neg^ cells **(Figure 4a)**. We found that mTORC1 activity was not uniformly distributed during quiescence but was strongly enriched in self-renewing *deltaA*:Gfp^neg/weak^ NSCs (61.0 ± 1.6%) compared to neurogenic *deltaA*:Gfp^high^ NSPCs (27.7 ± 2.4%) **(Figure 4b)**. This pattern suggests that mTORC1 activity during quiescence is not a generic feature of all NSPCs, but selectively highlights a large fraction of self-renewing NSCs at any given time.

**Figure 4.**
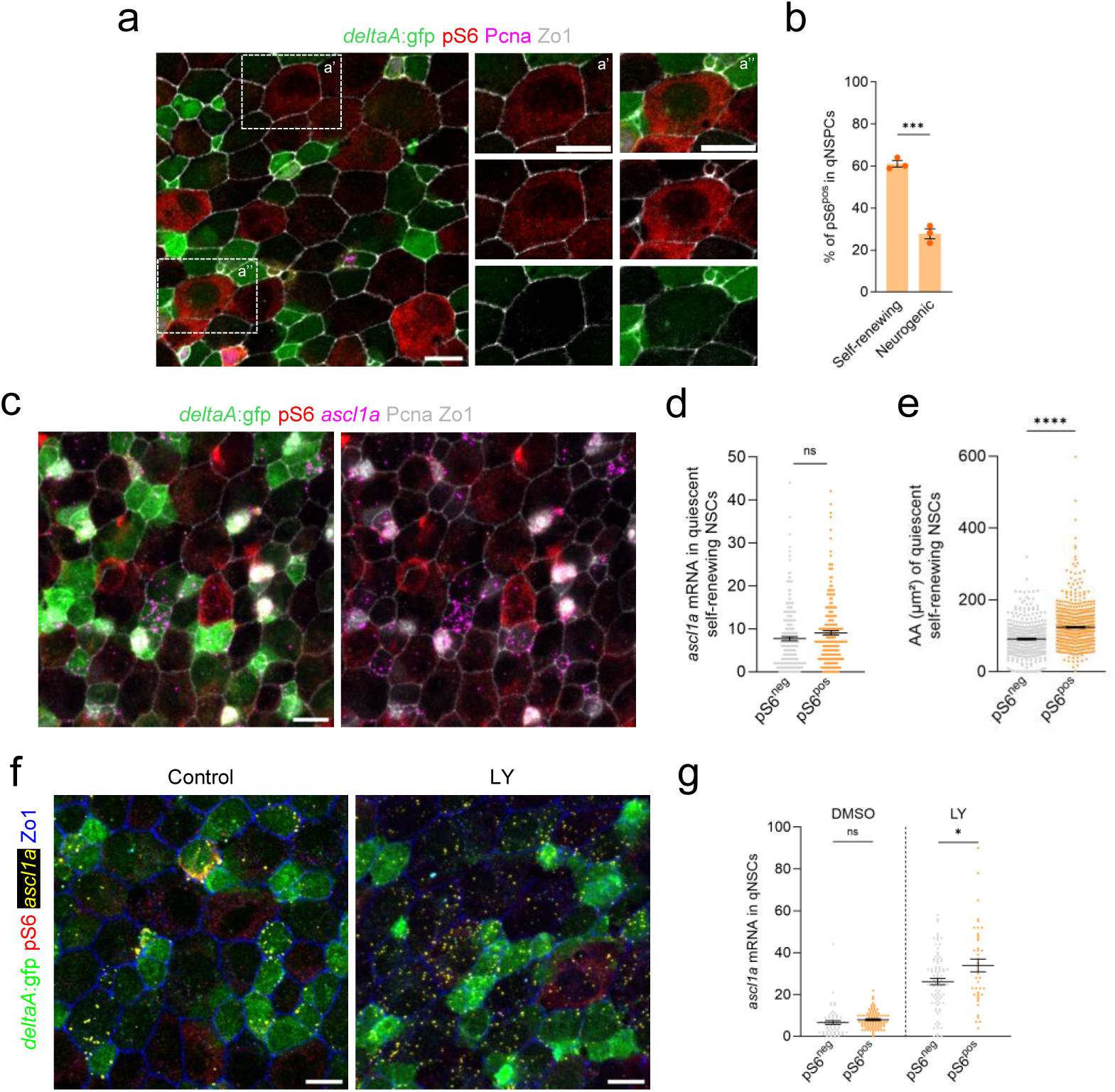
mTORC1 identifies a progression phase towards activation competence along the quiescence trajectory of self-renewing NSCs. **(a)** Adult Tg(*deltaA*:Gfp) pallia stained for pS6 (red), Pcna (magenta), Gfp (green), and Zo1 (white) showing pS6 expression in quiescent (Pcna^neg^) self-renewing NSCs (*deltaA*^neg^ (a’) and *deltaA*^weak^ (a”)). **(b)** Percentage of pS6^pos^ cells among quiescent self-renewing and neurogenic progenitors. **(c)** Expression of *ascl1a* mRNA (purple) revealed by smRNA-FISH on whole-mount Tg(*deltaA*:Gfp) adult pallia, together with IHC for pS6 (red), Gfp (green), and Pcna together with Zo1 (white). **(d)** *ascl1a* mRNA expression (number of puncta) in pS6^neg^ and pS6^pos^ quiescent self-renewing (*deltaA*^neg/weak^) NSCs. **(e)** Quantification of apical areas (AA, µm²) using Zo1 staining in pS6^neg^ and pS6^pos^ quiescent NSCs. **(f)** Whole-mount confocal images from adult Tg(*deltaA*:Gfp) fish, treated for 24h with DMSO or LY and analyzed by smRNA-FISH for *ascl1a* expression, coupled with IHC for Gfp (green), pS6 (red) and Zo1 (blue). **(g)** Quantification of *ascl1a* expression (number of puncta) in pS6^neg^ and pS6^pos^ quiescent self-renewing (*deltaA*^neg/weak^) NSCs upon 24h DMSO or LY treatment. NSC: neural stem cell. NSPCs: neural stem and progenitor cells. Data are shown as mean ± SEM from n=3 (b), n=2 (d, e) and n=1 (g) independent experiments. Two-tailed Mann–Whitney test (b) and unpaired t-test with Welch correction in (d, e, g). p-values : ****<0.0001; ***<0.001; **<0.01; *<0.05. Scale bars : 10 µm (a, c, f).

### mTORC1 activity signs the progressive acquisition of activation competence within the self-renewing NSC pool

Distinct quiescent substates with differing metabolic and activation potential, referred to as quiescence depths, have been previously identified in the adult zebrafish pallium (Morizet et al. 2025; Labusch et al. 2024). We therefore asked whether mTORC1 signaling within quiescent self-renewing NSCs is associated with progression along these substates. In mouse, high levels of the transcription factor Ascl1 mark NSCs engagement into activation, with its upregulation preceding and being required for cell-cycle entry (Andersen et al. 2014; Dulken et al. 2017; Rigo et al. 2025). *ascl1a* expression in Pcna^neg^ cells can also be used as a pre-activation marker in adult zebrafish NSCs (Morizet et al. 2025; Labusch et al. 2024). To test whether pS6^pos^ qNSCs are positioned in such a pre-activation state, we combined pS6 and Pcna IHC with *ascl1a* smRNA-FISH in adult Tg(*deltaA*:Gfp) pallia **(Figure 4c)**. As expected, qNSPCs expressed overall lower *ascl1a* levels than proliferating cells. In particular, within the self-renewing NSCs pool, pS6^neg^ and pS6^pos^ qNSCs showed only a modest difference in *ascl1a* expression (8.1 ± 0.6 in pS6^neg^ vs 9.8 ± 0.8 in pS6^pos^) **(Figure 4d)**. Thus, mTORC1 activity in self-renewing NSCs does not directly coincide with early transcriptional engagement into activation, suggesting that pS6 expression does not simply mark pre-activated cells but instead a different type of internal progression.

Because mTORC1 regulates anabolic growth and metabolism (He et al. 2025), we next examined whether pS6^pos^ qNSCs exhibit hallmarks of shallow or “primed” quiescence, a state described in several stem cell systems, and characterized by increased cell size, elevated mitochondrial metabolism, and heightened responsiveness to activation cues, distinguishing it from deeper states (van Velthoven et Rando 2019). We previously showed by long-term live imaging that self-renewing qNSCs gradually enlarge their apical area (AA) as they approach activation (Mancini et al. 2023). Using Zo1 to delineate apical cell contours in Tg(*deltaA*:Gfp) pallia, we found that quiescent pS6^pos^ self-renewing NSCs displayed significantly larger AA (123.6 ± 2.4 µm²) than their pS6^neg^ counterparts (90.7 ± 2.4 µm²) **(Figure 4e)**. This positions pS6^pos^ NSCs further along this intrinsic growth trajectory and temporally closer to activation.

To functionally test whether mTORC1 activity is associated with a quiescence phase where NSCs gain competency for activation, we examined the response of pS6^pos^ and pS6^neg^ self-renewing qNSC subsets to a forced push towards activation in vivo. We treated Tg(*deltaA*:Gfp) adult fish for 24h with the Notch inhibitor LY and monitored pre-activation and proliferation using *ascl1a* and Pcna **(Figure S4a, S4b)**. As previously described (Morizet et al. 2025), this treatment induced *ascl1a* expression (*deltaA*^neg/weak^ : 4.2 ± 0.1 vs 23.4 ± 0.5 mRNA puncta; *deltaA*^high^ : 4.9 ± 0.4 vs 14.8 ± 0.5 mRNA puncta in DMSO vs LY, respectively) **(Figure S4c)** yet without a significant increase in Pcna (*deltaA*^neg/weak^ : 1.2 ± 0.5 vs 3.4 ± 1.1; *deltaA*^high^ : 25.8 ± 2.9 vs 34.5 ± 5.8 cells in DMSO vs LY, respectively) **(Figure S4d)**, confirming the engagement of self-renewing NSCs (*deltaA*:Gfp^neg/weak^) towards activation within this treatment window. Following LY treatment, *ascl1a* expression increased in both populations but to a greater extent in pS6^pos^ self-renewing qNSCs (33.8 ± 3.0 mRNA puncta; 4.25-fold relative to DMSO) compared to pS6^neg^ qNSCs (26.2 ± 1.4 mRNA puncta; 4.0-fold relative to DMSO) (Figure 4i,j). This is consistent with mTORC1 activity marking qNSCs that are more responsive to activation cues and positioned closer to activation along the quiescence-to-activation trajectory **(Figure 4f, 4g)**. Thus, mTORC1 activity identifies qNSCs that are more responsive to activation cues and therefore positioned closer to activation along the quiescence-to-activation trajectory.

Together, our data show that mTORC1 activity does not coincide with pre-activation per se but instead marks the progression toward an activation-competent state within the self-renewing NSC pool. This state is characterized by enlarged apical size and enhanced responsiveness to activation signals. mTORC1 activity thus signs position along an intrinsic growth and priming trajectory that precedes division, with a substantial fraction of self-renewing NSCs residing within this phase at any given time, as inferred from the proportion of pS6^pos^ NSCs.

### mTORC1 inhibition biases the distribution of self-renewing NSCs along a conserved quiescence-activation trajectory without altering core transcriptional identity

To investigate the role of mTORC1 signaling in regulating adult quiescent NSC states, we performed single-cell RNA-sequencing (scRNA-seq) of adult pallial cells under control (DMSO) and mTORC1-inhibited (Rapamycin) conditions. Adult Tg(*sox2*:Gfp), in which Gfp labels NSPCs, were treated with either DMSO or Rapamycin for 24h or 72h, after which their pallia were dissected and cells dissociated. To enrich for the neurogenic lineage while preserving representation of downstream progeny, cells were FACS-sorted from each condition as a 50:50 mix of Gfp^pos^ and Gfp^neg^ cells prior to 10X Genomics library preparation **(Figure 5a)**. After quality filtering, 24,284 Rapamycin-treated cells and 19,647 control cells were retained for analysis across both timepoints and biological replicates **(Figure S5a, S5b).**

**Figure 5.**
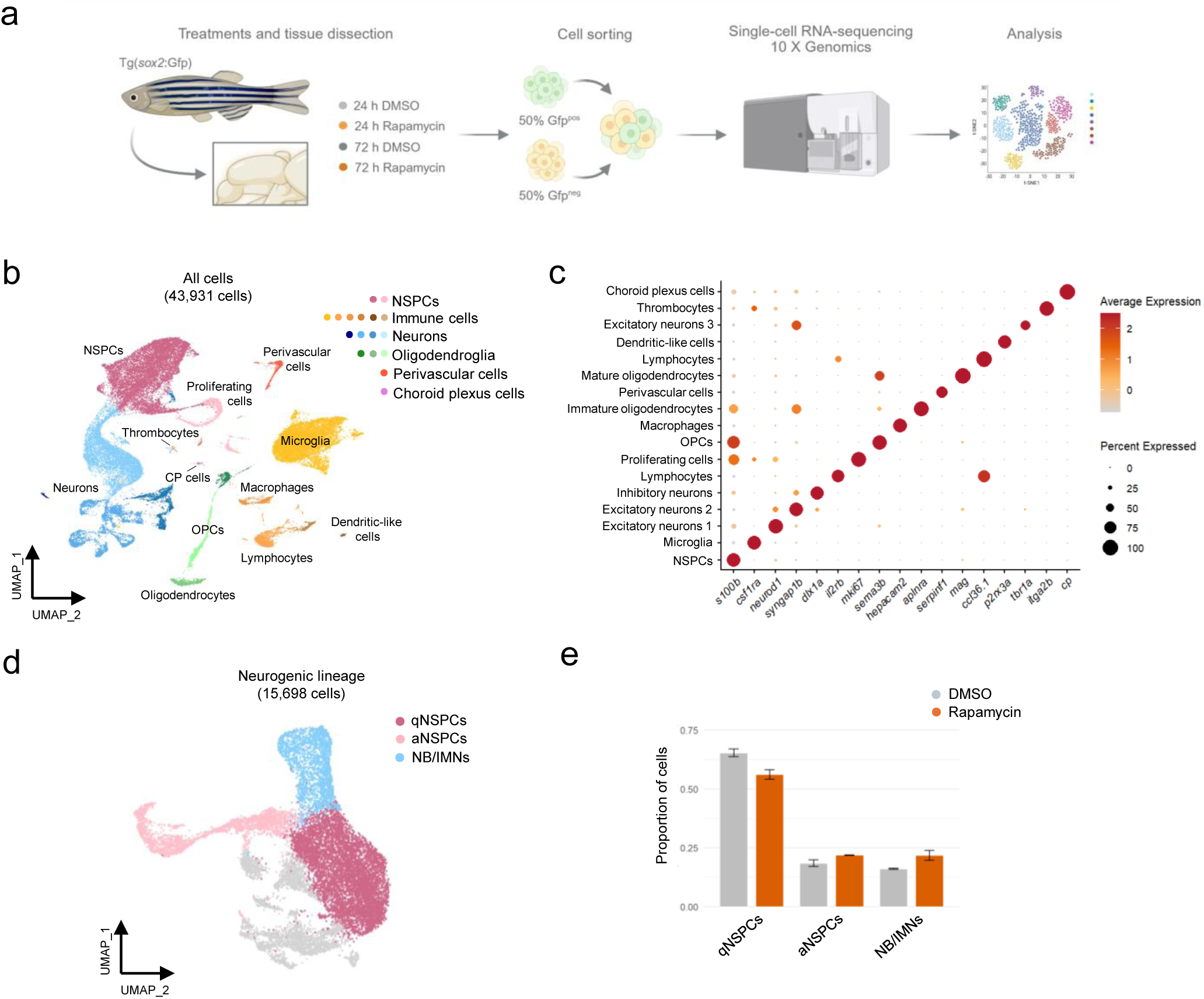
mTORC1 inhibition biases NSC state occupancy along the quiescence-to-activation trajectory. **(a)** Experimental design for scRNA-seq of adult Tg(*sox2*:Gfp) pallial cells after 24h and 72h DMSO or Rapamycin treatment (generated with BioRender). **(b)** UMAP embedding of all pallial cells across replicates and conditions, annotated by major cell populations. **(c)** Dot plot showing canonical marker genes’ expression defining major adult zebrafish pallial cell types. **(d)** UMAP of the neurogenic lineage (quiescent, activated NSPCs, NB/IMN), after subsetting from the full dataset. **(e)** Relative proportions of neurogenic clusters across conditions, showing redistribution of quiescent, activated NSPCs and NB/IMN upon Rapamycin treatment.

Integrated analysis revealed strong overlap in overall cell-state distributions across conditions and timepoints **(Figure S5c, S5d)**, indicating that short-term mTORC1 inhibition does not globally disrupt major cell-type identities. Unsupervised clustering identified 16 distinct pallial populations, well separated by their global transcriptome and annotated using canonical marker genes **(Figure S5e)**. These populations included NSPCs (*sox2*, *gfap*, *s100b*), 3 populations of pallial excitatory neurons at different maturation levels (*eomesa*, *neurod1*, *syngap1b*), 1 cluster of subpallial inhibitory neurons (*dlx1a*, *dlx5a*, *gad2*), as well as OPCs (*sema3b*) and oligodendrocytes (*aplnra*, *mag*). Several immune populations were recovered, including microglia (*csf1ra*, *mpeg1.1*), macrophages (*hepacam2*, *mpeg1.1*), 2 populations of lymphocytes (*ilr2b* and *ccl36.1* positive), thrombocytes (*ctfga*) and dendritic-like cells (*p2rx3a*). A cluster of perivascular cells (*serpinf1*) and choroid plexus cells (*cp*, *c4*) were also identified **(Figure 5b, 5c, Table S1)**.

We next focused on the neurogenic lineage, comprising qNSPCs, aNSPCs and neuroblasts (NBs)/immature neurons (IMNs). Subsetting yielded 9,386 Rapamycin-treated and 6,312 control cells **(Figure 5d)**, corresponding to clusters 6, 3, 1, 13, 11 (qNSPCs); 7, 16, 30 (aNSPCs) and 5 (NBs/IMNs) of the whole dataset **(Figure S5f)**. Control and Rapamycin-treated cells were broadly intermingled across the neurogenic lineage embedding, suggesting that short-term mTORC1 inhibition does not markedly alter overall lineage structure or cell-state distribution **(Figure S6a)**. We resolved 18 transcriptionally distinct clusters, corresponding to quiescent NSPCs (cl. 0, 1, 2, 4 and 5), activated NSPCs (cl. 6, 10, 13, 14) and neuroblasts (cl. 3, 8) **(Figure S6b)**, as well as spatially restricted NSPC populations consistent with pallial patterning domains (Morizet et al. 2024), including *gsx2*^pos^ cells (cl. 7, and 9) at the pallial-subpallial boundary and *nkx2.1*^pos^ of the ventral telencephalon (cl. 16) **(Figure S6b)**. Remaining clusters could not be clearly annotated. Among the pool of cells co-expressing NSPC markers (*s100b, sox2*, *gfap*), aNSPCs could be distinguished from qNSPCs based on the upregulation of cell cycle genes (*mki67*, *mcm2*, *pcna*) and neuroblasts based on neuronal specification genes (*eomesa*, *insm1a*, *neurod1*) **(Figure S6c)**. Quantification of cell-state proportions revealed a reproducible redistribution upon Rapamycin treatment **(Figure 5e)**, characterized by a relative depletion of quiescent NSPCs, specifically clusters 1, 2 and 5 **(Figure S6d)**, and a concomitant increase of aNSPCs and neuroblast states **(Figure 5e)**. This redistribution was consistent across both 24h to 72h treatments **(Figure S6e)**, suggesting an alteration in NSC state occupancy with a potential drift toward activation and/or commitment.

To more specifically assess the impact on self-renewing qNSCs, we sub-clustered the qNSPC compartment **(Figure 6a)**. Clusters enriched in self-renewing qNSCs (clusters 1, 2, and 5) were identified by highest expression of stem cell markers (ie. *mfge8a, hey1, cx43*) and lowest *deltaA* (*dla*) expression **(Figure S6f)**. The resulting UMAP embeddings showed good overlap between DMSO and Rapamycin **(Figure 6b)**. To match this UMAP with TORC1 activity, we derived an mTORC1 transcriptional activity module based on canonical mTORC1 downstream targets and projected it along the UMAP **(Figure 6c, Table S2).** Previous works have defined a continuum of NSC quiescence depth ranging from deep to shallow quiescence, followed by progressive activation, associated with coordinated metabolic, transcriptional, and cell cycle changes (Llorens-Bobadilla et al. 2015; Labusch et al. 2024; Dulken et al. 2017; Basak et al. 2018; Leeman et al. 2018; Jaehoon Shin et al. 2015; Morizet et al. 2025). Accordingly, the mTORC1 activity score increased progressively along an axis transitioning towards genes associated with preparedness to activation, e.g., encoding translational machinery (**Figure 6d**). This activation competence gradient followed the same conserved trajectory structure under DMSO and Rapamycin treatment. However, plotting cell proportions along this axis showed a subtle redistribution of cells towards cell states normally endowed with high mTORC1 scores in brains treated with Rapamycin vs DMSO-treated for 24h (**Figure 6e**). Together, these results demonstrate that mTORC1 inhibition does not rewire the transcriptional architecture of adult NSCs but instead biases their distribution along a conserved quiescence–activation continuum. Short-term inhibition transiently slows progression through this trajectory, without altering the underlying gene expression program. We conclude that mTORC1 signaling is integrated as a quantitative regulator of state progression rather than a determinant of cell identity.

**Figure 6.**
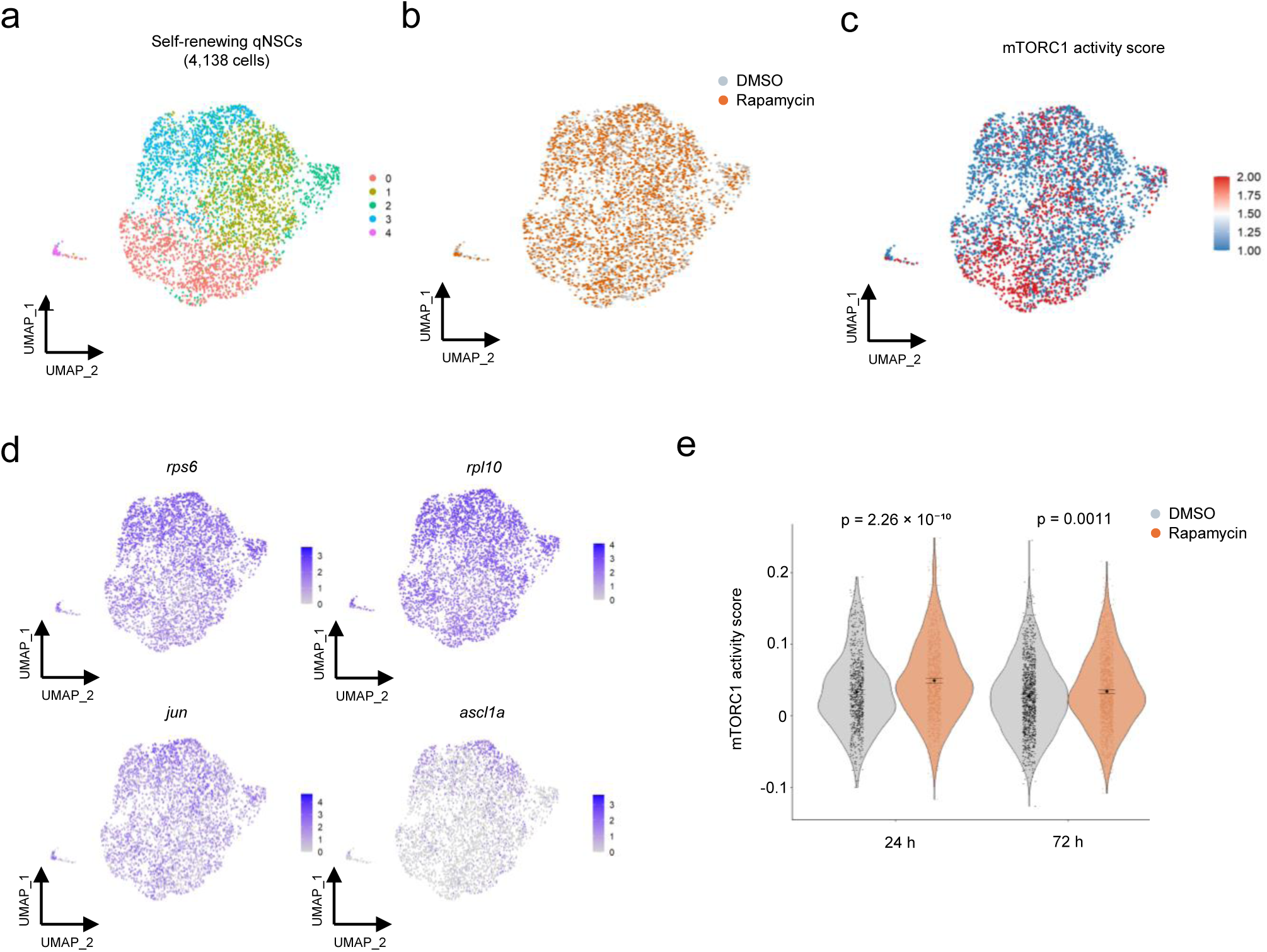
mTORC1 activity defines a conserved activation competence gradient in self-renewing quiescent NSCs. **(a)** UMAP of self-renewing quiescent NSCs identified within the neurogenic lineage. **(b)** UMAP embedding of self-renewing qNSCs across conditions. **(c)** mTORC1 transcriptional activity module score in self-renewing NSCs. **(d)** UMAP visualization of translational and activation-associated markers. **(e)** Cell density along mTORC1 activity score comparing conditions (DMSO: gray, Rapamycin: orange, after 24h (left) and 72h (right) of treatment). Black points indicate the mean, with error bars representing the 95% confidence interval. Statistical comparisons between treatments were performed using a two-sided Wilcoxon rank-sum test at the single-cell level, and p-values are indicated on the plots.

**Figure 7.**
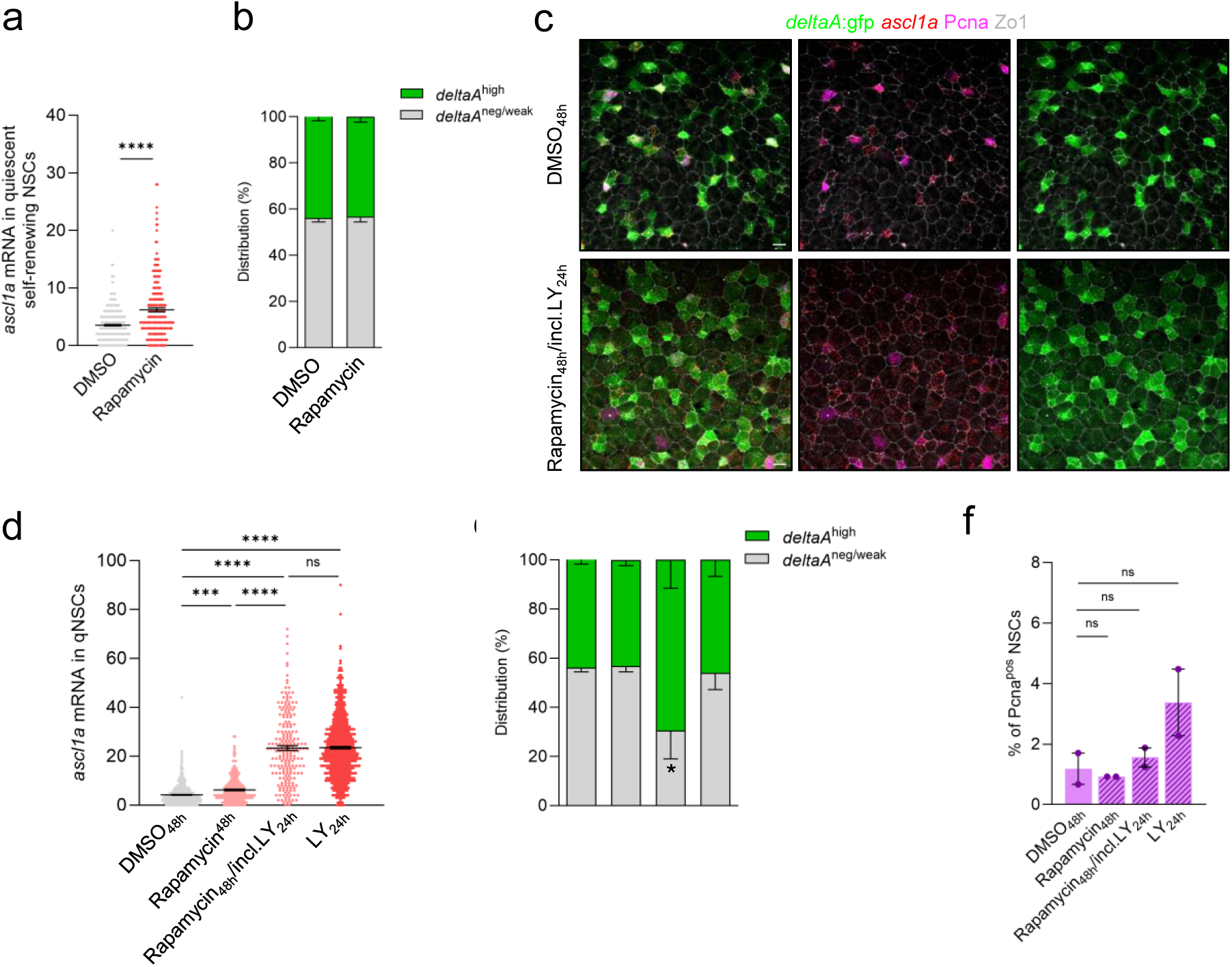
Effects of Rapamycin pre-treatment on NSC activation and lineage progression. **(a)** Quantification of *ascl1a* expression (number of puncta) in quiescent NSCs following 48h DMSO or Rapamycin treatment. **(b)** Distribution of *deltaA*:Gfp among NSPCs upon 48h DMSO or Rapamycin treatment. **(c)** Representative confocal images of *ascl1a* smRNA-FISH and Gfp (green), Pcna (purple) and Zo1 (white) IHC from adult Tg(*deltaA*:gfp) pallia after Rapamycin_48h_/incl.LY_24h_ treatment compared to DMSO controls. **(d)** Quantification of *ascl1a* levels (number of puncta) across treatment conditions (DMSO_48h_, Rapamycin_48h_, Rapamycin_48h_/incl.LY_24h_ and acute LY_24h_). **(e)** Distribution of *deltaA*^neg/weak^ and *deltaA*^high^ NSPCs in DMSO_48h_, Rapamycin_48h_, Rapamycin_48h_/incl.LY_24h_ and LY_24h_ treatments. **(f)** Percentage of Pcna^pos^ NSPCs across DMSO_48h_, Rapamycin_48h_, Rapamycin_48h_/incl.LY_24h_ and LY_24h_ conditions within self-renewing NSCs (*deltaA*^neg/weak^). Data shown as mean ± SEM from n=2 independent experiments. Unpaired t-test with Welch correction in (a, d) and two-tailed Mann–Whitney test (f). p-values : ****<0,0001; ***<0,001; **<0,01 ; *<0,05. Scale bars : 10µm.

### mTORC1 inhibition in shallow quiescence promotes differentiation-prone activation responses in self-renewing NSCs

Proportional shifts in scRNA-seq populations can hint at changes in tissue composition, but differences in dissociation efficiency and cell survival across cell types limit their direct interpretation. We therefore used these changes only as initial indicators and validated all conclusions with tissue-level by IHC. To test these interpretations in situ, we first examined the effect of 48h Rapamycin alone on the transition of quiescent self-renewing NSCs towards activation. Adult Tg(*deltaA*:Gfp) fish were treated with Rapamycin and expression of the pre-activation marker *ascl1a* was analyzed with smRNA-FISH **(Figure 6a, 6c)**. Rapamycin modestly increased *ascl1a* expression in self-renewing NSCs (mean number of puncta: *deltaA*^neg/low^ : 6.2 ± 0.3 in Rapamycin vs 3.5 ± 0.1 in DMSO) **(Figure 6c)**, while *deltaA* subtype proportions remained unchanged across self-renewing (*deltaA*^neg/weak^ : 56.7 ± 2.3% upon Rapamycin) and neurogenic populations (*deltaA*^high^ : 43.3 ± 2.3%) compared to controls (*deltaA*^neg/weak^ : 56.1 ± 1.7% and *deltaA*^high^: 43.92 ± 1.7%) **(Figure 6b)**. These results indicate that mTORC1 inhibition shifts some quiescent self-renewing NSCs toward a pre-activation state without altering lineage composition under basal conditions.

To determine whether this shift affects subsequent fate engagement, we applied a modified two-step paradigm in which adult Tg(*deltaA*:Gfp) fish were treated with Rapamycin for 48h and Notch inhibition with LY was applied during the final 24h of Rapamycin treatment. Under these conditions, we expect to test the fate decision of cells pushed towards quiescence exit by LY in a context where mTORC1 has been blocked during a longer period. Under these conditions, self-renewing NSCs, whether or not receiving a Rapamycin treatment prior to LY, similarly displayed a marked induction of *ascl1a* (*deltaA*^neg/weak^ : 23.3 ± 14.5 in Rapamycin_48h_/incl.LY_24h_ and 23.5 ± 0.5 in LY vs 3.7 ± 0.2 in DMSO) **(Figure 6d)**. However, Rapamycin_48h_/incl.LY_24h_ treatment also increased the proportion of *deltaA*^high^ progenitors (69.5 ± 11.6% compare 43.9 ± 1.7% in DMSO) compared to *deltaA*^neg/weak^ NSCs (30.5 ± 11.6% in Rapamycin_48h_/incl.LY_24h_ vs 56.1 ± 1.7 in DMSO), an effect not observed with LY alone **(Figure 6e)**. Thus, mTORC1 inhibition modifies the responsiveness of self-renewing NSCs to activation cues and promotes their transition toward a neurogenic state. Notably, across all conditions (DMSO, Rapamycin, Rapamycin_48h_/incl.LY_24h_, and LY treatments), the proportion of Pcna^pos^ cells among self-renewing NSCs remained unchanged **(Figure 6f)**, indicating that these effects occur without a corresponding increase in proliferative entry. Together, these results show that mTORC1 inhibition increases the propensity of qNSCs to engage activation-associated transcriptional programs and lineage progression, without promoting cell-cycle entry. Consistent with our scRNA-seq analyses, these findings support a model in which mTORC1 activity restrains premature progression along the quiescence-to-activation continuum and preserves the coupling between activation and self-renewal.

## DISCUSSION

Our work uncovers an unexpected heterogeneity related to mTORC1 signaling within the quiescent NSC population under physiological homeostatic conditions. Previous models proposed that adult NSCs exhibit very low mTORC1 activity, which gradually rises as cells approach activation and peaks in proliferating progenitors (Nieto-González et al. 2019; Zhou et al. 2018). By contrast, we show that a large fraction of long-term self-renewing NSCs displays detectable mTORC1 activity during quiescence, a feature that is largely absent from neurogenic NSPCs. We further show that mTORC1-positive self-renewing NSCs exhibit features associated with a graded continuum towards the acquisition of activation competence, characterized by increased apical size, enhanced metabolic activity, and a transcriptional state biased toward activation. These features are reminiscent of the G_alert_ state in MuSCs (Rodgers et al. 2014) and the LRIG1⁺ primed NSC pool in the adult mouse SVZ (Marqués-Torrejón et al. 2021). However, whereas alert or primed states have typically been associated with injury, inflammation, or aging, our results demonstrate that mTORC1 similarly marks such activation competence acquisition in quiescence under physiological conditions. This suggests that mTORC1-dependent priming represents a general mechanism by which stem cells regulate activation competence within a quiescence continuum, rather than a state exclusively induced by stress.

NSC maintenance in the adult brain is crucially dependent on quiescence (Ehm et al. 2010; Imayoshi et al. 2010; Ables et al. 2010). In this context, our findings offer new insights into the regulation of quiescence within the self-renewing NSC population of the zebrafish adult pallial niche, the most upstream NSC of the lineage maintaining life-long stemness (Mancini et al. 2023; Than-Trong et al. 2020). First, our data reinforce the idea that quiescence is a heterogeneous and dynamically regulated continuum rather than a fixed identity. Self-renewing NSCs at any given time differ in their mTORC1 activity, and the association of mTORC1 with a transcriptome and metabolic state poised for activation suggests that it reflects progression along an activation competence continuum rather than a discrete sub-lineage. Additionally, the association of mTORC1 with NSCs of large AA, together with the documented progressive growth of AA during quiescence, supports the idea that mTORC1-positive NSCs represent an intermediate state along a continuous trajectory toward activation, rather than a separate quiescence depth program. Our pharmacological manipulations of mTORC1, analyzed at transcriptomic and tissue levels, show that mTORC1 activity inhibition does not abolish this trajectory but shifts NSCs along it, accelerating entry into early activation while impairing proper maintenance of stemness. Thus, we propose that mTORC1 activity supports a metastable quiescence sub-state in which NSCs progressively acquire activation competence while maintaining long-term self-renewal potential. Disruption of this balance biases NSCs toward differentiation-prone activation, suggesting that controlled progression through this state is essential to preserve stem cell identity. At any given time, nearly 60% of self-renewing NSCs in quiescence can be found in the mTORC1^pos^ state, suggesting that the acquisition of activation competence is an extended process that accounts for roughly half of the duration of quiescence on average. Together, our results position mTORC1 as a key regulator of quiescence dynamics and activation competence, consistent with emerging views that stem cell quiescence is actively maintained and tuned by the intensity and integration of multiple signaling inputs, rather than being a static state (Urbán et al. 2019; Cheung et Rando 2013).

In mammalian systems, apical surface organization and cell–cell contact geometry have been linked to stem cell behavior and responsiveness to niche signals (Shaya et al. 2017; Mirzadeh et al. 2008), suggesting that morphological features may reflect differences in activation state or responsiveness to cell-cycle entry cues. In our study, we found that mTORC1 activity in quiescent self-renewing NCS correlates with increased AA and increased expression of translation machinery genes, two hallmarks of a metabolically primed state. Whether these features depend on mTORC1 and are functionally relevant for this NSC state remains to be tested. Nevertheless, these observations suggest that mTORC1 may coordinate cellular growth and metabolic readiness with the early steps of cell cycle re-entry. This is consistent with the established role of mTORC1 as an integrator of nutrient and growth factor signaling, which promotes both cell growth and G1/S cell cycle progression. In the context of quiescence, mTORC1 activity may act as a molecular hub that links cellular growth with activation competence, rather than simply proliferation onset. Along a more speculative line, we note that a large AA correlates on average with more neighboring cells, which may impact signaling received by the local niche. Thus, AA growth during shallow quiescence may also rewire how NSCs integrate niche-derived signals, affecting their sensitivity to activation cues. While this hypothesis remains to be directly tested, it would offer a mechanistic framework linking metabolic growth to niche-sensing behavior during acquisition of activation competence.

The upstream signals regulating mTORC1 activity during quiescence in adult NSCs remain to be identified. Insights from other stem cell systems suggest potential candidates: in MuSCs, Gli transcription factors have been shown to modulate mTORC1 signaling, thereby influencing activation and fate decisions (Peng et al. 2023; Brun et al. 2022). In the context of adult NSCs, recent work has identified Myc as a key regulator promoting the transition from shallow quiescence to activation. mTORC1 regulates c-Myc through multiple mechanisms (He et al. 2025) and in particular, Myc and mTORC1 cooperate in cancer to control protein synthesis and cell growth (Liu et al. 2017; Pourdehnad et al. 2013). Whether similar crosstalk operates in adult NSCs to regulate activation competence within quiescence remains an important question for future investigation.

mTORC1 acts at multiple steps along the quiescence-to-activation sequence, regulating progression towards activation competence during a long quiescence phase, supporting early activation, and enabling full NSC activation and progression through the cell cycle. Although our data demonstrate these sequential requirements, we could not directly resolve a graded mTORC1 signaling continuum at the single cell resolution. pS6 levels are heterogeneous among qNSCs and increase in cycling NSPCs, but do not resolve discrete stepwise thresholds that could reflect changes in signaling strength. However, our results provide the first integrated experimental evidence that mTORC1 acts coordinately across multiple stages of the NSC quiescence-to-activation continuum, a model that had previously been inferred only indirectly from independent studies examining its role in NSC activation, proliferation and fate decisions (Hartman et al. 2013; Zhou et al. 2018; Baser et al. 2019; Rodgers et al. 2014, 2017).

## METHODS

### Ethics

All procedures involving zebrafish were conducted in compliance with the European Directive 2010/63/EU on the protection of animals used for scientific purposes. Experimental protocols were approved by the competent authorities of the Paris region (Department 483) under institutional animal care and use agreements (facility authorization number C75-15-22).

### Zebrafish lines and husbandry

Zebrafish (*Danio rerio*) were maintained under standard laboratory conditions in a recirculating aquatic system. Fish were housed in 3.5-L tanks at a maximum density of five animals per liter. Water temperature was maintained at 28.5 °C with a pH of 7.4, under a 14 h light / 10 h dark cycle. Larvae were fed rotifers three times daily until 14 days post-fertilization, after which they were transitioned to a commercial dry diet (GEMMA Micro, Skretting). Adult fish aged 2.5 to 4 months were used for all experiments, irrespective of sex. The following lines were employed: Tg(*gfap*:Gfp)^mi2001^ (Bernardos et Raymond 2006), Tg(*gfap*:dTomato) (Satou et al. 2012), Tg(*deltaA*:Gfp) (Madelaine et Blader 2011), Tg(*sox2*:Gfp) (Jimann Shin et al. 2014) and AB wild-type fish.

### Euthanasia

For tissue collection, fish were euthanized by immersion in ice-cold water (1–2 °C) for 10 min, in accordance with institutional guidelines and under specific authorization from the French Ministry of Higher Education, Research and Innovation.

### In vitro cultures

#### 2D NSC cultures

Neural stem cell (NSC) cultures were established from the adult zebrafish pallium as previously described in Thetiot et al. 2025 (Thetiot et al. 2025). Briefly, pallial tissues from ∼20 adult zebrafish were microdissected, pooled, and dissociated into single cells by enzymatic (FACSMax) and mechanical procedures. Cells were enriched for neural stem and progenitor populations by Percoll density gradient centrifugation (22% v/v). The resulting cell suspension was typically distributed into two wells of a 24-well plate. For 2D cultures, cells were plated onto plates sequentially coated with poly-D-lysine (10 µg.mL⁻¹) and laminin (10 µg.mL⁻¹) at a density corresponding to ∼20 pallia per well. Cells were maintained in a defined serum-free medium consisting of DMEM/F12 supplemented with B-27 (1×), N2 (1×), insulin (2 µg.mL⁻¹), glucose (0.3% w/v), and antibiotic–antimycotic (1×). To promote NSC proliferation, the medium was supplemented with human FGF and EGF (20 ng.mL⁻¹ each). Cultures were maintained at 30 °C in a humidified incubator with 5% CO₂. After an initial 48 h adhesion period, non-adherent cells were collected, centrifuged, and replated onto the same wells to maximize recovery. Thereafter, 75% of the medium was renewed every 2–3 days until cell passaging.

#### 3D spheroids generation

For 3D cultures, enriched NSCs were seeded in 96-well plates (U-bottom) under non-adherent conditions in the same complete medium at a density of 1000 cells per well, allowing the formation of free-floating neurosphere-like spheroids. Spheroids were maintained at 30 °C with 5% CO₂ and fed every 2–3 days by 75% medium replacement.

### Pharmacological treatments

Pharmacological treatments were performed both in vivo and in vitro. Notch signaling was inhibited using the γ-secretase inhibitor LY-411575 (LY; Sigma-Aldrich, cat# SML0506-25MG), prepared as a 10 mM stock solution in DMSO, aliquoted, and stored at −20 °C. mTORC1 activity was inhibited using Rapamycin (Sigma-Aldrich, 553210-5MG), prepared as a 1 mM stock solution in DMSO, aliquoted, and stored at −20 °C. For in vivo treatments, fish were transferred to fresh tanks at a density of one animal per 50 mL of system water and exposed to LY-411575 at a final concentration of 10 µM or to Rapamycin at 0.2 µM. Control groups received an equivalent concentration of DMSO. Treatments were carried out for 24 h, 48h or 72h; with daily renewal of treatment solution. Fish were fed for 1h prior to each renewal. For in vitro experiments, NSC cultures were treated with LY-411575 (5 µM) or Rapamycin (20 nM) at the indicated time points, with equivalent concentrations of DMSO used as controls.

### Tissue preparation for immunohistochemistry and smRNA-FISH

Adult zebrafish were euthanized as described above, and brains were dissected in ice-cold PBS. Tissues were fixed in 4% paraformaldehyde (PFA) for 4 h at room temperature (RT) or overnight at 4°C with gentle agitation. After fixation, tissues were washed in PBS containing 0.1% Tween-20 (PBS-T) prior to dehydration through a graded methanol concentrations (25%, 50%, 75%, and 100% methanol in PBS-T), 5 min each incubation at RT, with gentle agitation. Samples were stored in 100% methanol at −20 °C until use.

### Immunohistochemistry

Prior to staining, tissues were rehydrated through a graded reverse methanol series (75%/50%/25% methanol in PBS-T), with 5 min incubations at RT under gentle agitation. Samples were then incubated in pigment removal solution (3% H_2_O_2_, 5% formamide, 0.5X SSC prepared in H2O) for 15 min under illumination, followed by 5 min wash in PBS-T at RT.

Antigen retrieval was performed in HistoVT One solution (Nacalai Tesque) at 65 °C for 1 h, followed by three 10 min washes in PBS-T. Tissues were subsequently incubated in blocking solution (4% goat serum, 0.1% Triton X-100, 0.1% DMSO in PBS) for at least 3 h at RT.

Primary antibodies were diluted in blocking solution and incubated overnight at 4 °C. The following primary antibodies were used : Chicken anti-GFP (1:500; Aves Labs, cat# GFP-1020), Mouse anti-Sox2 (Abcam, cat# ab171380), Mouse anti-Zo1 (clone ZO1-1A12; 1:200; ThermoFisher Scientific, cat# 33-9100), Mouse anti-Tomm20 (1:250; Santa Cruz Biotechnology, sc-17764); Mouse anti-Pcna (1:250; Santa Cruz Biotechnology, cat# sc-56), Rabbit anti-Sox2 (1:500; Abcam, cat# ab97959), Rabbit anti-pS6 (1:200; Ozyme, cat# 2211S); Rabbit anti-p4EBP1 (1:200; Ozyme, cat# 2855S), Rabbit anti-S6 (1:200; Ozyme, cat# 2317S), Rat anti-RFP (clone 5F8, 1:250, ChromoTek, cat# 5f8-100).

After three 10 min washes in PBS-T, samples were incubated with secondary antibodies diluted in blocking buffer under the same conditions. Secondary antibodies included: Goat anti-Chicken IgG(H+L), Alexa Fluor 488 Conjugated (1:1000; ThermoFisher Scientific, cat#A-11039), Goat anti-Mouse IgG1, Alexa Fluor 546 conjugated (1:1000; ThermoFisher Scientific, cat# 10082322), Goat anti-mouse IgG1, DyLight™ 405 (1:1000; Biolegend, cat# 409109), Goat anti-Mouse IgG2a, Alexa Fluor 633 conjugated (1:1000; ThermoFisher Scientific, cat# A-21136), Goat anti-Rabbit IgG(H+L), Alexa Fluor 546 Conjugated (1:1000; ThermoFisher Scientific, cat# A-11010), Goat anti-Rabbit IgG(H+L), Alexa Fluor 488 Conjugated (1:1000; ThermoFisher Scientific, cat# A11008), Goat anti-Rat IgG(H+L), Alexa Fluor 546 Conjugated (1:1000; ThermoFisher Scientific, cat# 10143662). Samples were then washed three times 10 min in PBS-T at RT under gentle agitation.

When incubated, DAPI (1:10000, Merck, cat# d8417) diluted in blocking solution was applied for 30 min following immunostaining and followed by three 10 min washes in PBS-T.

Dissected pallia were mounted in PBS-T using 0.5-mm spacers and sealed with Valap (a mixture of petrolatum, paraffin, and lanolin). 3D spheroids were processed using the same immunostaining protocol and mounted in PBS-T using depression slides (Thermo Fisher Scientific, cat# 11875930).

### Multiplex RNA in situ hybridization (HiPlex)

Multiplex fluorescent in situ hybridization was performed using the RNAscope® HiPlex v2 platform (Bio-Techne) following the manufacturer’s instructions, with adaptations for zebrafish whole-mount brains. All reagents were equilibrated at RT prior to use. Tissue were incubated in probe diluent buffer for a 3 h pre-hybridization step at 40°C with gentle agitation, followed by overnight hybridization at 40 °C with target probes (diluted 1:50 and pre-warmed for 10 min) with gentle agitation. The following probe was used : RNAscope® HiPlex Probe - *Danio rerio* (achaete-scute family bHLH transcription factor 1a (*ascl1a*); Bio-Techne, cat# 583661-T2).

Samples were then washed three times 10 min in RNAscope wash buffer at RT under gentle agitation. Signal amplification was performed through sequential incubations with HiPlex amplification reagents (Amp1–Amp3), each for 30 min at 40 °C with agitation, with three 10 min washes in RNAscope wash buffer between each step. Fluorescent detection was achieved by incubation with fluorophore reagents (T1–T3) for 15 min at 40 °C with agitation. Immunohistochemistry was subsequently performed following completion of the hybridization protocol.

### Single Cell RNA Sequencing

#### Sample preparation (dissociation and cell sorting)

Brains were dissected from adult Tg(*sox2*:Gfp) zebrafish in ice-cold 1× PBS and kept on ice throughout the procedure. For each condition, approximately 20 fish were pooled per replicate without sex selection. Two independent biological replicates per condition were generated on separate days; within each replicate, all four conditions (DMSO and Rapamycin treatments at 24 h and 72 h) were processed in parallel.

Tissues were enzymatically dissociated using FACSMax dissociation solution (30 °C, 5 min), followed by gentle mechanical trituration through a 40 μm cell strainer to obtain a single-cell suspension. Cells were centrifuged at 400 × g for 5 min at 4 °C and resuspended in ice-cold 1× PBS supplemented with 0.04% BSA. Cell viability was assessed using 7-AAD staining.

Cell suspensions were sorted using a FACSAria III (70 μm nozzle, low-pressure 4-way sort mode) to minimize mechanical stress, and approximately 30,000 viable cells per condition were collected in a final volume of ∼30 μL. Sorting gates were defined based on forward and side scatter to exclude debris and dead cells. GFP fluorescence from the Tg(*sox2*:Gfp) reporter line was used to separate Gfp^pos^ and Gfp^neg^ fractions, which were then recombined in a 1:1 ratio prior to library preparation. This strategy enabled enrichment of neural stem and progenitor cells while preserving representation of downstream and non-neurogenic populations, thereby maintaining the overall cellular diversity of the tissue.

#### Library preparation and sequencing

Single-cell suspensions were processed to generate cDNA libraries using the Chromium Single Cell 3′ Reagent Kit (10x Genomics, version 3.1), following the manufacturer’s protocol. Briefly, approximately 20,000 cells were loaded per channel with an expected recovery of 12,000 cells. All subsequent steps, including reverse transcription, cDNA amplification, and library construction, were performed according to the manufacturer’s instructions. Libraries were indexed using a Dual Index kit (10x Genomics) and pooled prior to sequencing. Sequencing was carried out on an Illumina platform (NovaSeqX) using paired-end reads configured according to 10x Genomics recommendations. Raw base call files were converted to FASTQ format and processed using the Cell Ranger pipeline (10x Genomics, version 7.2) with GRCz11 fasta and V4.3.2.gtf from Lawson lab zebrafish transcriptome annotation to generate gene expression matrices.

#### Data processing and quality control

Output from Cell Ranger was analyzed using the Seurat package (version 5.1.0) in R (version 4.3.0). Low-quality cells were excluded based on the following criteria: cells with fewer than 2000 detected genes and/or a proportion of mitochondrial gene expression exceeding 5% were removed. Additional filtering thresholds were adjusted as necessary to account for dataset-specific characteristics. To identify and exclude potential doublets, computational doublet detection was performed using scDblFinder (version 1.16.0). The expected doublet rate was estimated based on cell loading density and recommendations from 10x Genomics.

#### Normalization, dimensionality reduction, and clustering

After quality control, all experimental conditions (DMSO 24 h, DMSO 72 h, Rapamycin 24 h and Rapamycin 72 h) were re-extracted from raw counts and merged into one large Seurat object. Gene expression data were normalized using the `NormalizeData` function (scale factor = 10,000). Highly variable genes were identified using `FindVariableFeatures` (nfeatures = 1,000). Data were then scaled prior to principal component analysis (PCA), performed using the `RunPCA` function. Integration were performed using harmony (1.2.1) on sample to take account of inter sample variability. Dimensionality reduction was carried out using Uniform Manifold Approximation and Projection (UMAP) via the `RunUMAP` function on integrated reduction, based on the first 30 principal components selected using the elbow plot. Nearest-neighbor graphs were constructed using `FindNeighbors` (number of PCs = 30) on integrated reduction, and clusters were identified using the `FindClusters` function (resolution = 1.0).

#### Identification and characterization of clusters

Cluster-specific marker genes were identified using the `FindAllMarkers` function from Seurat, using the Wilcoxon statistical framework. Differentially expressed genes were selected based on thresholds of adjusted *P* value < 0.05 and log fold-change > 0.25. Cell populations were annotated based on the expression of known marker genes derived from published literature and publicly available datasets (Morizet et al. 2024; Cosacak et al. 2019; Pandey et al. 2023; Anneser et al. 2024). Cluster identities were assigned manually by comparing the expression patterns of canonical markers across clusters.

After initial clustering, neurogenic populations were identified based on the expression of established marker genes, including quiescent neural stem/progenitor cells (NSPCs), proliferating cells, and neuroblasts. Quiescent NSPCs were characterized by high expression of stem cell markers (*sox2*, *s100b*) and low expression of differentiation markers (*neurod1*, *eomesa, elavl3*). Proliferating cells were defined by the expression of cell-cycle–associated genes (e.g., mcm2, *pcna, mki67*) together with NSPC markers, while neuroblasts were identified by the expression of early neuronal differentiation markers (e.g., *neurod1*, *eomesa, elavl3*). These neurogenic populations were subsetted from the full dataset and re-analyzed from raw counts. Data normalization and scaling, followed by dimensionality reduction, integration were performed using CCAintegration. Clustering were performed to refine population structure within the neurogenic compartment. To further resolve quiescent self-renewing NSC heterogeneity, clusters corresponding to cells with highest levels of *mfge8a* and low levels of *dla* were retained.

This stepwise approach resulted in multiple Seurat objects corresponding to different levels of resolution, including the full dataset, the neurogenic compartment and enriched quiescent NSCs. These analyses were performed across integrated experimental conditions (DMSO 24 h, DMSO 72 h, Rapamycin 24 h and Rapamycin 72 h) as well as per timepoints (24 h vs 72 h treatments).

#### Differential gene expression analysis

Differential gene expression (DEGs) analysis between clusters or experimental conditions was performed using the `FindMarkers` or `FindAllMarkers` functions in Seurat. Genes were considered significantly differentially expressed if they met the criteria of adjusted *P* value < 0.05 and avg_log2fc > 0.25. Additional filtering thresholds were applied where appropriate.

#### mTORC1 transcriptional module score

mTORC1 pathway activity was quantified using the HALLMARK_MTORC1_SIGNALING gene set from MSigDB (v2026.1). Gene lists were imported from .gmt files and parsed in R. Because the dataset was generated in zebrafish, human genes were converted to zebrafish orthologs using the babelgene package. Only orthologs detected in the dataset were retained to define the final module. The complete gene list is provided in Supplementary Table S2. mTORC1 Module score was computed at single-cell resolution using the `AddModuleScore` function from Seurat, which calculates the average expression of the gene set normalized against control gene sets with matched expression levels. The resulting score was added to cell metadata and used for downstream visualization on UMAP embeddings.

### Image acquisition

Fluorescence imaging was performed using LSM700 confocal microscope or LSM710 confocal microscope equipped with a 40× oil immersion objective (Plan-Apochromat 40×/1.3 NA). Image acquisition was controlled using ZEN (Blue or Black editions). Acquisition parameters, including laser power (0.5–8%), detector gain (600–900), and z-step size, were optimized for each experiment and kept constant across samples within the same experimental set. Images were acquired at 16-bit depth with a resolution of 1024 × 1024 pixels, 2X averaging per line and zoom factors ranging from 0.5× to 1.5×.

### Image analysis

#### 3D spheroids

Quantification of cytoplasmic (*gfap*:dTomato, S6, pS6) and nuclear (Sox2, Pcna, DAPI) signals in 3D spheroids was performed using Fiji. For each spheroid, a central optical plane was selected and combined with the immediately adjacent z-planes (two above and two below) to generate a projected image approximating the thickness of a single nucleus. Cells were identified based on Sox2 or DAPI staining, and signal quantification was then performed with manual assignment of individual cells. Nuclei that were below a defined size threshold were excluded from the analysis to avoid segmentation artifacts.

#### Adult zebrafish pallium

Image analysis was performed using a custom napari plugin (FishFeats (Letort et al. 2026)) developed for 3D analysis of adult pallial surface cells. This tool enables the quantification of junctional (Zo1), cytoplasmic (Gfp, S6, pS6, p4EBP1, Tomm20), nuclear (Pcna, Sox2), and smRNA-FISH (*ascl1a*) signals. Junctional staining (Zo1) was projected into 2D using a local maximum projection and segmented with EpySeg (Aigouy et al. 2020). Segmentations were manually corrected when necessary using dedicated tools within the plugin. Apical cell area was extracted from the corrected Zo1 segmentation. smRNA-FISH signals were detected using Big-FISH with automated threshold estimation. Detected spots were initially assigned to cells based on their projection onto the apical surface. Spots located outside of segmented apical regions were left unassigned. All assignments were subsequently manually curated to ensure accurate attribution of each RNA punctum to its most probable cell. Spots detected in multiple channels, likely representing imaging artifacts, were excluded from analysis. The number of RNA puncta per cell was then quantified. Cytoplasmic immunostaining intensities were analyzed by assigning cells to user-defined categories using the “ClassifyCells” function implemented in FishFeats.

### Statistical analysis

Statistical analyses were performed using GraphPad Prism (GraphPad Software, version 9.5.1). Data are presented as mean ± SEM. Unless otherwise specified, each data point represents an independent biological replicate (individual animal or independent culture). No statistical methods were used to predetermine sample size. Comparisons between two groups were performed using two-tailed unpaired tests: Student’s t-test when variances were comparable, Welch’s t-test when variances were unequal, and Mann–Whitney test for small sample sizes where normality could not be assumed. All tests were two-sided. For experiments with more than two conditions, pairwise comparisons were conducted using unpaired t-tests with Welch’s correction, as indicated in the figure legends.

Imaging-based quantifications of cell proportions were averaged per biological replicate prior to statistical testing. In contrast, measurements of apical area and *ascl1a* smRNA-FISH signal (number of puncta) were quantified at the single-cell level, with statistical analyses performed on pooled cell values across replicates. For scRNA-seq analyses, changes in cell proportions were interpreted descriptively and validated experimentally. Statistical analyses were performed in R. Comparisons of mTORC1 activity scores between treatment groups were conducted using a two-sided Wilcoxon rank-sum test at the single-cell level, as implemented in the wilcox.test function. P-values were reported as indicated in the figures. A p-value < 0.05 was considered statistically significant (*p* < 0.05, p < 0.01, *p* < 0.001, p < 0.0001).

## AUTHOR CONTRIBUTIONS

Conceptualization : M.T., L.B-C.; Methodology: M.T., L.T., D.M.; Writing original draft: M.T; Review and editing of original draft: M.T., L.B-C; Supervision: L.B-C., Project administration: L.B-C.; Funding acquisition: L.B-C., M.T. All authors have read and agreed to the published version of the manuscript.

## ACKNOWLEDGMENTS

We thank Sébastien Bedu and Nathan Guibert for expert fish care, members of the L.B-C team for their support and discussions. We acknowledge the Center for Translational Science (CRT)- Cytometry and Biomarkers Unit of Technology and Service (CB UTechS) at Institut Pasteur for support in conducting this study. This work was funded by the ANR (Labex Revive – ANR-10-LABX-0073), the Fondation pour la Recherche Médicale (EQU202203014636), the European Research Council (ERC SyG PEPS 101071786), CNRS and Institut Pasteur (Explore Program) to L.B-C. Melina Thetiot was supported by a Pasteur-Roux-Cantarini fellowship from Institut Pasteur and a LabEx Revive fellowship.

