## Supplemental figures and legends for "mTORC1 supports progression toward activation competence in quiescent adult neural stem cells"

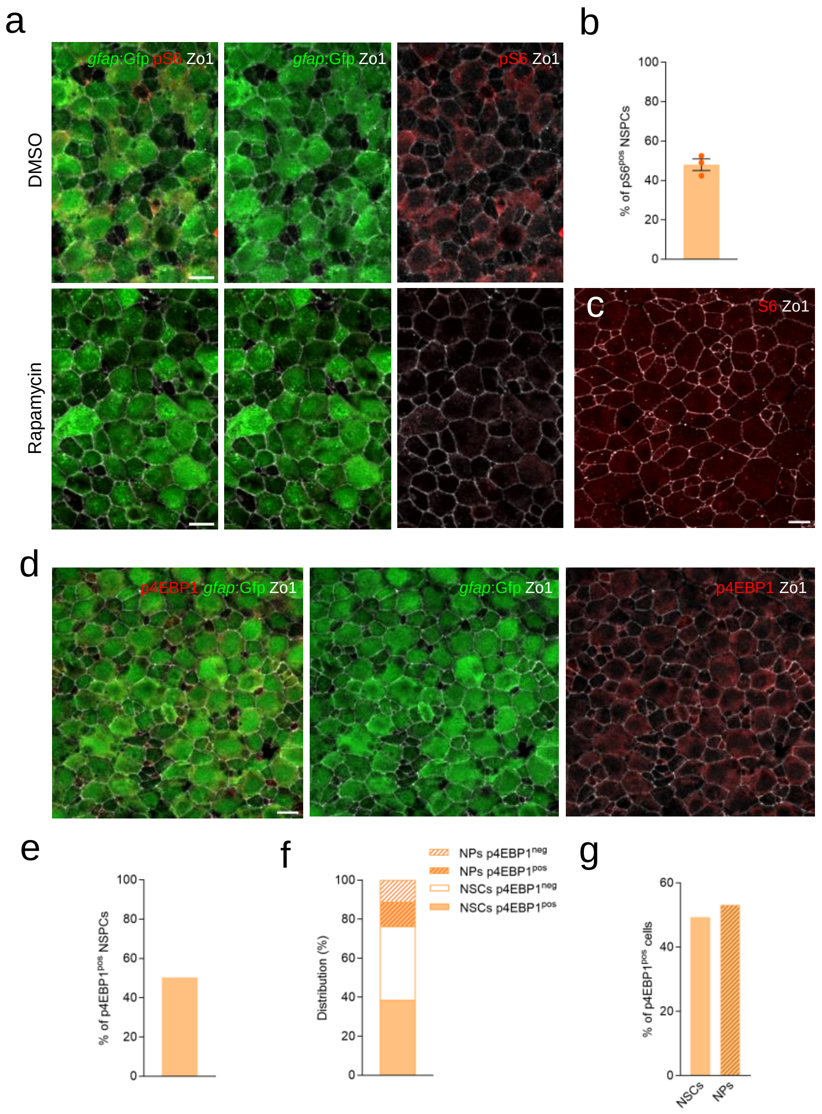
a
gfap:Gfp pS6 Zo1
gfap:Gfp Zo1
pS6 Zo1
DMSO
Rapamycin
b
% of pS6pos NSPCs
c
S6 Zo1
d
p4EBP1 gfap:Gfp Zo1
gfap:Gfp Zo1
p4EBP1 Zo1
e
% of p4EBP1pos NSPCs
f
Distribution (%)
NPs p4EBP1neg
NPs p4EBP1pos
NSCs p4EBP1neg
NSCs p4EBP1pos
g
% of p4EBP1pos cells
NSCs
NPs

Figure S1

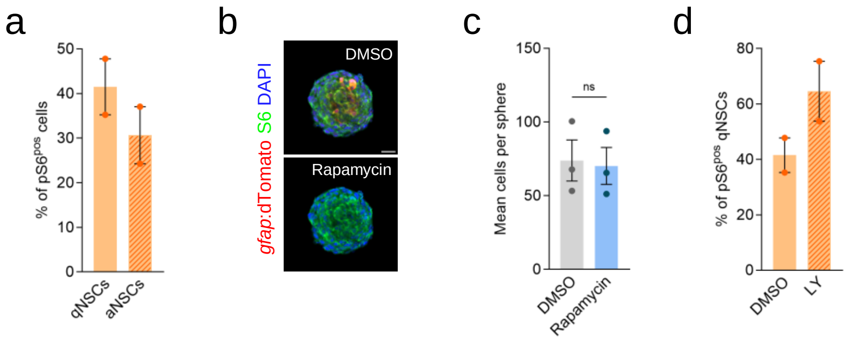

a
50
40
30
20
10
0
% of pS6pos cells
qNSCs
aNSCs
b
DMSO
Rapamycin
gfap:dTomato S6 DAPI
c
150
100
50
0
Mean cells per sphere
ns
DMSO
Rapamycin
d
80
60
40
20
0
% of pS6pos qNSCs
DMSO
LY


Figure S2

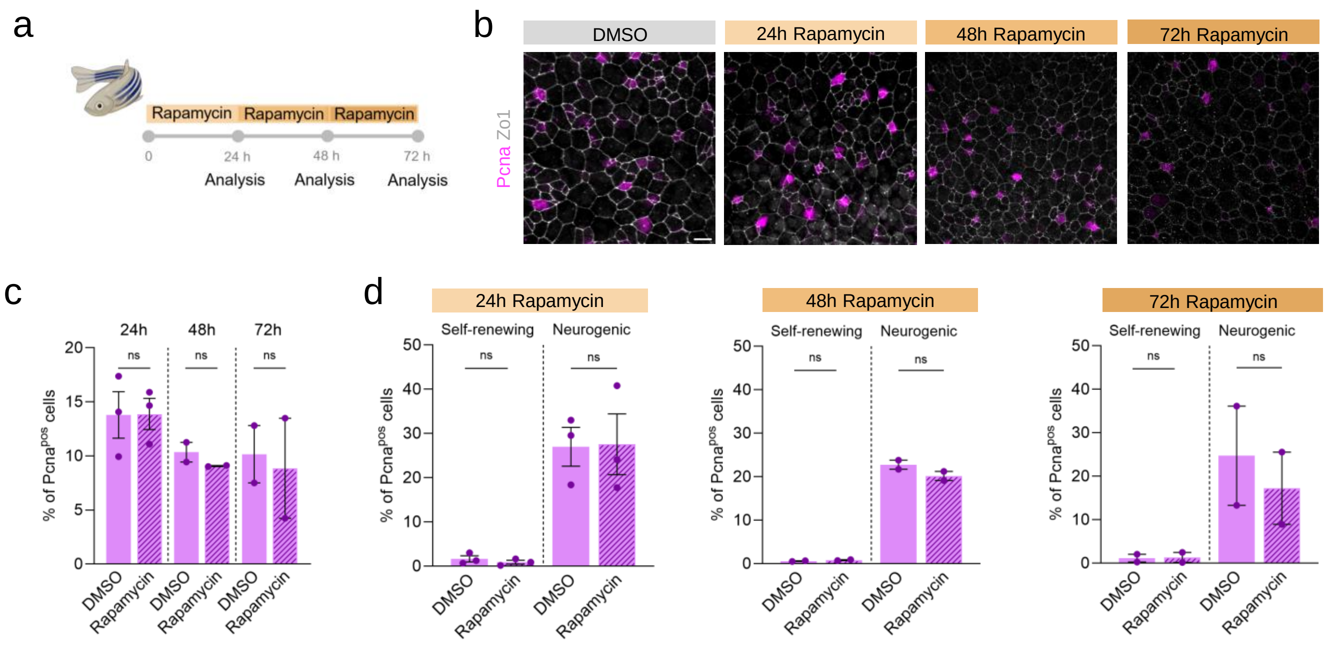
a
Rapamycin
Rapamycin
Rapamycin
0
24 h
48 h
72 h
Analysis
Analysis
Analysis
b
DMSO
24h Rapamycin
48h Rapamycin
72h Rapamycin
Pcna Zo1
c
24h
48h
72h
ns
% of Pcna$^{pos}$ cells
DMSO
Rapamycin
d
24h Rapamycin
48h Rapamycin
72h Rapamycin
Self-renewing
Neurogenic
ns
% of Pcna$^{pos}$ cells
DMSO
Rapamycin

Figure S3

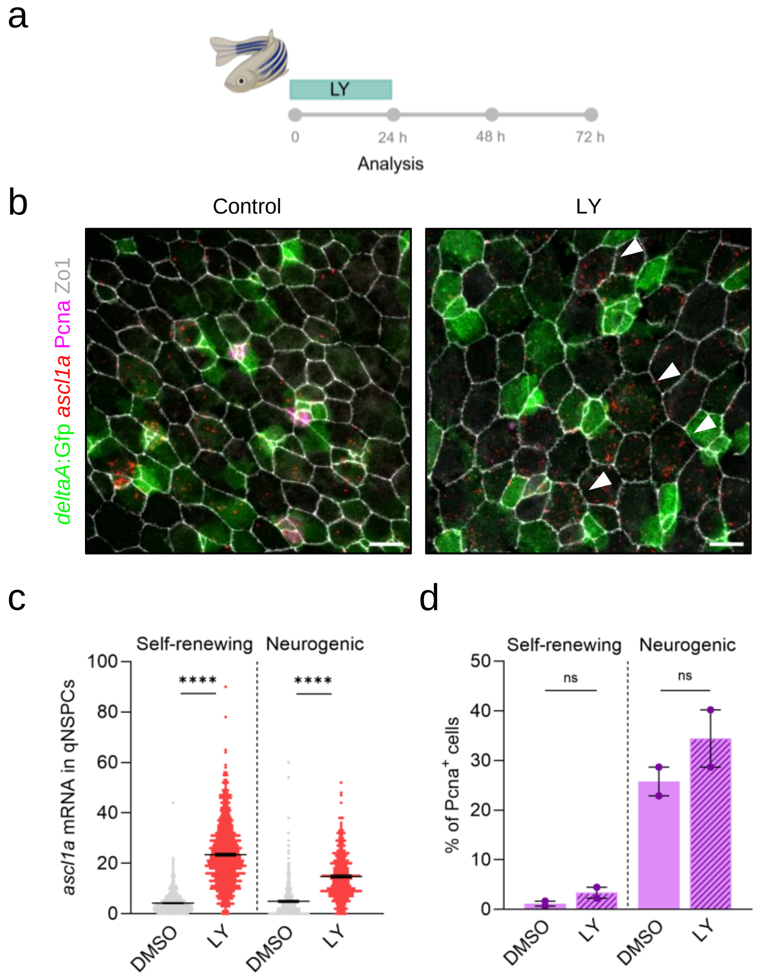
a
LY
0
24 h
48 h
72 h
Analysis
b
Control
LY
deltaA:Gfp ascl1a Pcna Zo1
c
Self-renewing
Neurogenic
ascl1a mRNA in qNSPCs
****
****
DMSO
LY
DMSO
LY
d
Self-renewing
Neurogenic
% of Pcna+ cells
ns
ns
DMSO
LY
DMSO
LY

Figure S4

a

| Library name | Number of fish | Sex | Number of cells |
|---|---|---|---|
| R1_DMSO_1D | 20 | M+F | 4138 |
| R2_DMSO_1D | 20 | M+F | 4436 |
| R1_DMSO_3D | 20 | M+F | 4847 |
| R2_DMSO_3D | 20 | M+F | 6226 |
| R1_RAPA_1D | 20 | M+F | 5532 |
| R2_RAPA_1D | 20 | M+F | 6054 |
| R1_RAPA_3D | 19 | M+F | 5780 |
| R2_RAPA_3D | 20 | M+F | 6918 |

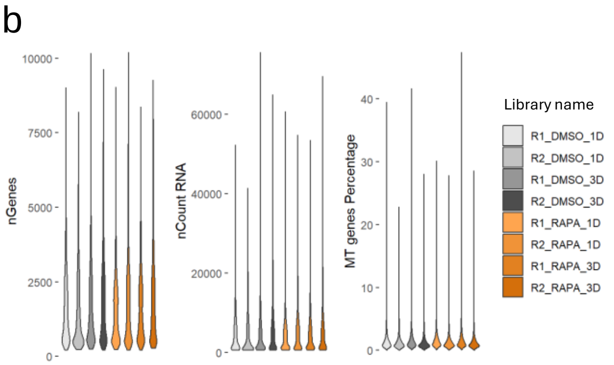


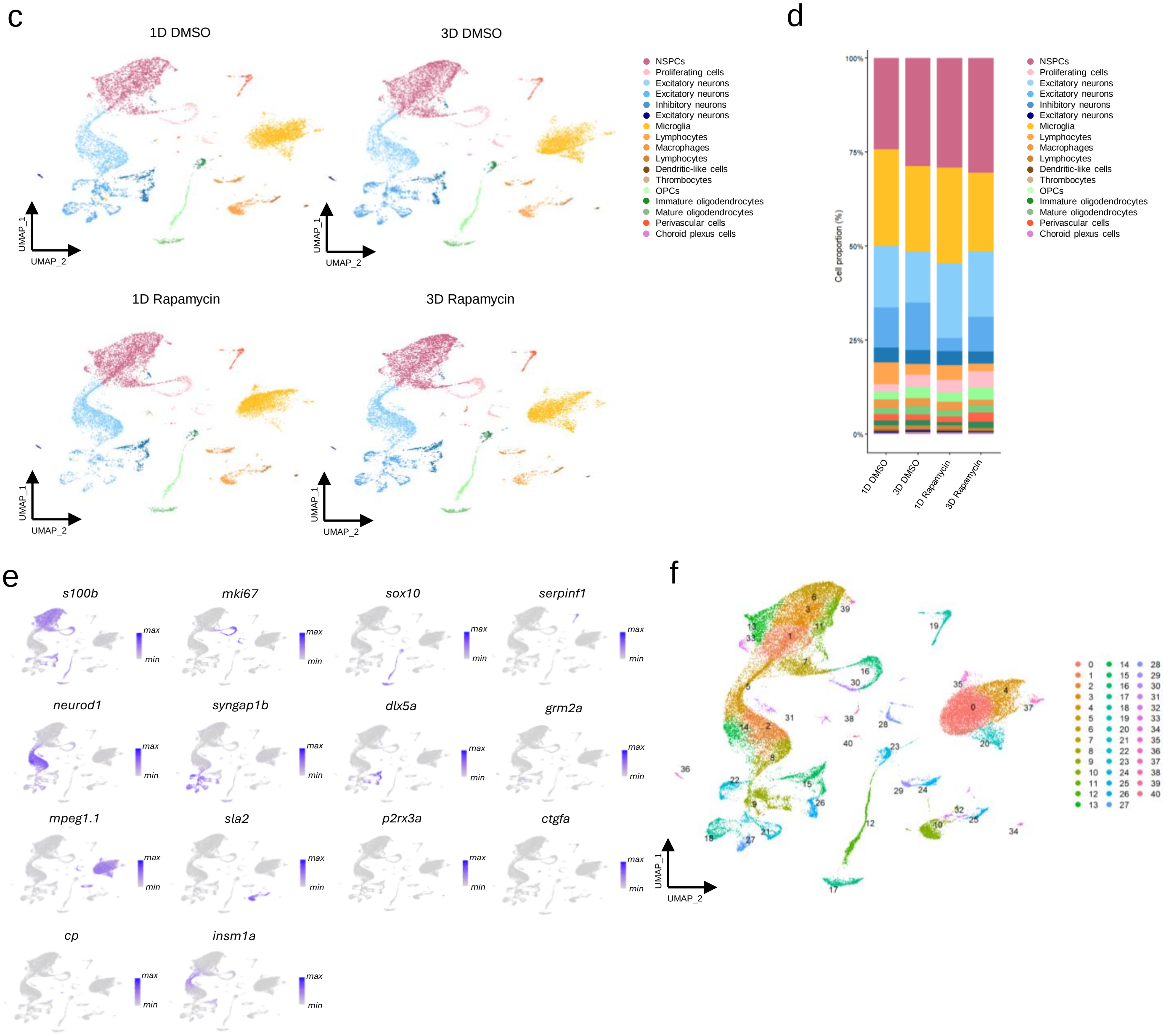


Figure S5

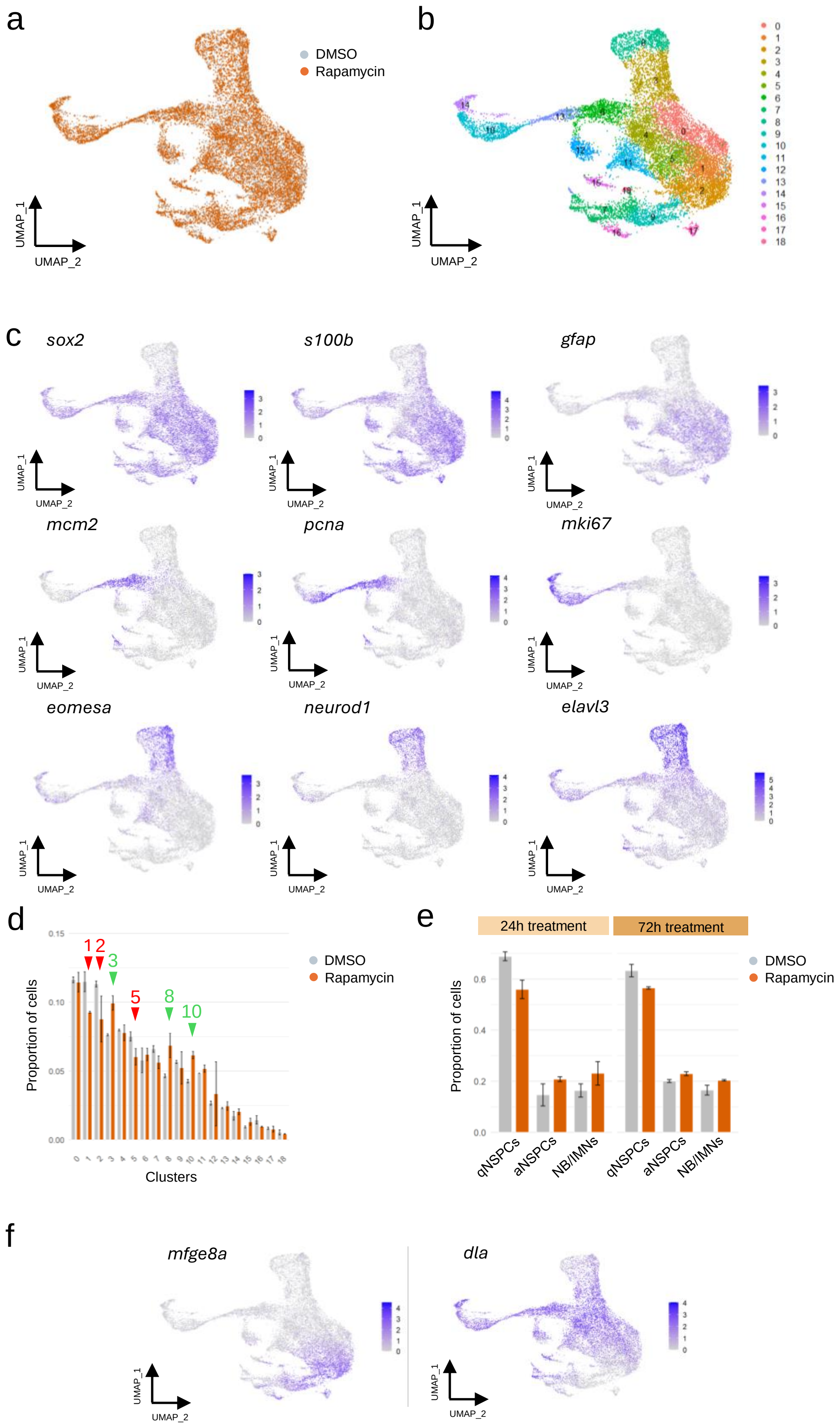
a
DMSO
Rapamycin
UMAP_1
UMAP_2
b
UMAP_1
UMAP_2
c
sox2
s100b
gfap
mcm2
pcna
mki67
eomesa
neurod1
elavl3
d
Proportion of cells
Clusters
DMSO
Rapamycin
e
24h treatment
72h treatment
DMSO
Rapamycin
Proportion of cells
qNSPCs
aNSPCs
NB/IMNs
f
mfge8a
dla

Figure S6

### SUPPLEMENTARY FIGURE LEGENDS

**Figure S1. mTORC1 activity in NSCs is reflected by post-translational phosphorylation of downstream targets.**

**(a)** Whole-mount IHC from 3-month-old Tg(*gfap*:Gfp) adult fish treated with DMSO or Rapamycin for 24 h, showing reduced pS6 signal (red) in NSPCs following Rapamycin treatment. Individual NSPCs are delineated by Zo1 (white). **(b)** Quantification of the proportion of pS6$^{pos}$ cells among total NSPCs. **(c)** Representative whole-mount IHC from a 3-month-old Tg(*gfap:*Gfp) adult zebrafish pallium, stained for total S6 (red) and Zo1 (white). **(d)** Representative IHC from a Tg(*gfap*:Gfp) adult zebrafish pallium at 3-month, stained for phosphorylated 4EBP1 (p4EBP1, red), Gfp (green), and Zo1 (white). **(e)** Quantification of p4EBP1$^{pos}$ and p4EBP1$^{neg}$ among total pallial NSPCs. **(f)** Distribution of p4EBP1$^{pos}$ across *gfap*$^{pos}$ NSCs and *gfap*$^{neg}$ NPs, manually assigned (full and stripped orange, respectively). **(g)** Percentage of p4EBP1$^{pos}$ cells among *gfap*$^{pos}$ NSCs and *gfap*$^{neg}$ NPs. NSPC: neural stem and progenitor cells. Data shown as mean ± SEM from n=3 (b) and n=1 (e, f, g) independent experiments. Scale bars : 10 µm (a, c, d).

**Figure S2. Validation of mTORC1 inhibition in vitro.**

**(a)** Percentage of pS6$^{pos}$ qNSCs (Pcna$^{neg}$) and aNSCs (Pcna$^{pos}$) (full and stripped orange, respectively) in 3D spheroids at 3 DIV. **(b)** Representative confocal images of 3D spheroids from Tg(*gfap*:dTomato) fish showing homogeneous S6 expression across NSCs in both DMSO and Rapamycin-treated conditions, stained for S6 (green), dTomato (red), and DAPI (blue). **(c)** Mean normalized number of NSCs within the spheroids in DMSO and Rapamycin-treated conditions. **(d)** Percentage of pS6$^{pos}$ qNSCs (Pcna$^{neg}$) in DMSO and LY-treated conditions (full and stripped orange, respectively). Data shown as mean ± SEM from n=2 (a, d) and n=3 (c) independent experiments. Two-tailed Mann–Whitney test. p-values : ****<0.0001; ***<0.001; **<0.01; *<0.05. Scale bar: 10 µm (b).

**Figure S3. In vivo characterization of NSPCs responses to Rapamycin.**

**(a)** Schematic representation of Rapamycin treatments procedure. **(b)** Confocal image of the dorsal pallium from Tg(*deltaA*:Gfp) adult fish treated in vivo for 24 h, 48h and 72h with DMSO or Rapamycin, stained for Gfp (green), Pcna (magenta), and Zo1 (white). **(c)** Percentage of Pcna$^{pos}$ NSPCs after 24h, 48h and 72h DMSO or Rapamycin treatments. **(d)** Percentage of Pcna$^{pos}$ NSPCs after 24h, 48h and 72h DMSO or Rapamycin treatments in *deltaA*$^{neg/weak}$ self-renewing NSCs and *deltaA*$^{high}$ neurogenic NSPCs. Scale bars : 10 µm (a). Data shown as mean ± SEM from n = 2 to 3 independent experiments. Two-tailed Mann–Whitney test (c, d). p-values : ****<0.0001; ***<0.001; **<0.01; *<0.05.

**Figure S4. Dynamics of NSPCs response to pushed activation.**

**(a)** Schematic representation of the LY411575 (LY) treatment procedure. **(b)** Confocal images from Tg(*deltaA*:Gfp) treated with LY411575 (LY) for 24h to inhibit Notch signaling showing *ascl1a* (smRNA-FISH, red) and immunolabeled Gfp (green), Pcna (purple) and Zo1 (white). White arrowheads point to increased *ascl1a* expression among qNSPCs. **(c)** Quantification of *ascl1a* mRNA (number of puncta) in quiescent self-renewing (*deltaA*$^{neg/weak}$) NSCs and neurogenic (*deltaA*$^{high}$) NSPCs upon DMSO or LY treatments. **(d)** Percentage of Pcna$^{pos}$ cells across *deltaA*:Gfp subpopulations (self-renewing and neurogenic) upon 24h LY treatment compared to control (DMSO). NSPCs: neural stem and progenitor cells. Data shown as mean ± SEM from n = 2 independent experiments. Unpaired t-test with Welch correction in (c) and two-tailed Mann–Whitney test (d). p-values : ****<0,0001; ***<0,001; **<0,01; *<0,05. Scale bars : 10 µm (b).

**Figure S5. scRNA-seq dataset overview and quality control.**

**(a)** Summary of 10x Genomics libraries and sequencing metrics. **(b)** Violin plots of the number of genes, counts and the percentage of genes per cells across libraries. **(c)** Integrated UMAP of all cells showing robust overlap of control and Rapamycin conditions and both timepoints. **(d)** Relative proportions of all cell populations in the zebrafish pallium, across conditions. **(e)** Feature plots showing canonical marker expression across all pallial cell types. **(f)** UMAP of the neurogenic lineage showing clustering.

**Figure S6. Structure of the neurogenic lineage and transcriptional stability under mTORC1 inhibition.**

**(a)** Integrated embedding of the neurogenic lineage across treatment conditions. **(b)** UMAP of the neurogenic lineage. **(c)** Feature plots of key markers distinguishing NSPCs and neuroblasts/immature neurons. **(d)** Cluster-level composition changes upon Rapamycin treatment; arrowheads indicate depletion (green) and enrichment (red). **(e)** Comparison of neurogenic lineage composition (cell proportions between clusters) between 24 h and 72 h treatments under DMSO and Rapamycin treatment. **(f)** Markers used to select qNSC clusters enriched in self-renewing NSCs (high expression of *mfge8a*, low expression of *dla*). NSPC: neural stem and progenitor cells. NB/IMNs : neuroblasts/immature neurons.
